# Elflow: Automated Quality Control and Self-Supervised Representation Learning for High-Dimensional Cytometry

**DOI:** 10.64898/2026.11.17.767309

**Authors:** Yogesh Kalakoti, Maura Gasparetto, Sachin Kumar Gupta, Vaibhav Mittal, Abhishek Jha

## Abstract

High-dimensional flow and mass cytometry are foundational technologies for clinical immunophenotyping and hematological diagnostics, yet computational data analysis remains bottlenecked by operator-subjective manual gating. While computational tools have emerged, they remain fragmented across isolated tasks such as quality control or discrete clustering, leaving an unmet need for integrated, modular workflows that resolve continuous biological variation without manual supervision. Here, we introduce **Elflow**, a modular, self-supervised framework that links automated quality control, single-cell representation learning, and patient-level modeling without manual gating at inference. Elflow couples automated machine-learning quality control with Variance-Invariance-Covariance Regularization (VICReg) to learn continuous, batch-resilient single-cell representations without negative pairs or hand-labeled training sets. Across four diverse public benchmarks, Elflow adapts to distinct biological panels and clinical objectives. On 22-channel spectral flow cytometry (OMIP-030), self-supervised embeddings preserve fine functional T-cell gradations and outperform classical baselines in unsupervised clustering, while an anchor-guided extension accurately reproduces canonical manual gating hierarchies. On hand-curated CyTOF panels (Levine-13), linear PCA matches deep models, establishing an empirical boundary condition where linear methods suffice. Across multi-center clinical cohorts (FlowCAP-II), Elflow representations achieve high diagnostic sensitivity for acute myeloid leukemia across heterogeneous instruments despite severe class imbalance. In longitudinal immune monitoring (FlowCAP-IV), patient mean embeddings and micro-cluster differential histograms preserve rare antigen-specific responses, yielding significant AIDS-free survival stratification. By uniting automated quality control, self-supervised representation learning, and multi-scale patient aggregation, Elflow provides a modular, reproducibly benchmarked framework for high-parameter and multi-center translational cytometry.

## 1. Introduction

High-dimensional single-cell cytometry, encompassing poly-chromatic flow cytometry, full-spectrum spectral flow cytometry, and mass cytometry (CyTOF), is an indispensable pillar of modern immunology, hematology, and clinical oncology [1, 2]. By measuring 10 to over 40 surface and intracellular protein markers on millions of individual cells per patient sample, cytometry enables deep phenotypic dissection of immune cell subsets, clinical diagnosis of hematological malignancies, and immune monitoring during clinical trials [3, 4].

Despite profound advances in cytometer instrumentation and fluorophore chemistry, computational data analysis remains a formidable bottleneck. Historically, clinical analysis has relied on *manual hierarchical gating*, the sequential drawing of two-dimensional polygonal boundaries across pairs of markers by human analysts. Manual gating is not only labor-intensive and slow, but inherently operator-subjective. Multi-center studies conducted by the Human Immunology Project Consortium (HIPC) and the FlowCAP consortium have revealed inter-operator and inter-center coefficients of variation exceeding 20% to 30% on identical sample aliquots [5–7]. Furthermore, manual gating restricts discovery to preconceived, axis-aligned 2D projections, rendering it structurally incapable of detecting unforeseen phenotypes or mapping continuous immunological gradients.

To overcome these human limitations, a wide array of computational tools has been developed over the past decade [8, 9]. However, the current computational ecosystem remains fundamentally fragmented, with few integrated workflows connecting individual analysis stages. Existing unsupervised methods typically target only isolated stages of the workflow:

a. *Automated Quality Control (QC)*: Tools such as flowAI [10], flowClean [11], and PeacoQC [12] identify acquisition anomalies and instrument perturbations, but do not perform cell phenotyping or clinical prediction.
b. *Unsupervised Clustering*: Widely adopted algorithms such as FlowSOM [13], PhenoGraph [14], Citrus [15], and Cytofkit [16] automate cell subset grouping, but quantize continuous phenotypic variation into rigid, discrete meta-clusters. In doing so, they discard intra-cluster heterogeneity, struggle to resolve continuous functional expression patterns (such as chemokine receptor polarizations in helper T-cell subsets or fine activation/exhaustion gradients), and require arbitrary pre-specification of cluster counts.
c. *Dimensionality Reduction*: Non-linear projections such as *t*-SNE and UMAP [17] provide intuitive two-dimensional visualizations, but do not provide quantitative, batch-invariant latent representations suitable for cross-cohort clinical modeling.

Consequently, immunologists and clinical trialists are forced to cobble together disjointed software scripts, creating work-flow friction, compounding error propagation, and leaving downstream clinical predictions vulnerable to batch variation and optical spillover.

Furthermore, adapting contemporary deep representation learning frameworks directly to high-dimensional cytometry poses fundamental algorithmic challenges. Deep generative models such as scVI [18] were architected for sparse, discrete single-cell RNA-sequencing counts; when adapted to continuous cytometry intensities, their reconstruction objectives tend to be dominated by abundant resting cell types at the expense of rare, functionally pivotal populations. Standard self-supervised contrastive frameworks (e.g., SimCLR [19]) rely on treating distinct cells as “negative pairs” to be pushed apart in latent space. This negative-pair assumption directly conflicts with cytometric biology, where millions of cells across patients share near-identical baseline states (e.g., resting lymphocytes); forcing such cells apart degrades true biological topology. Finally, in deep 20-to 40-parameter panels, flat neural encoders suffer from *lineage dominance*: high-variance bimodal lineage markers (e.g., CD4 vs. CD8 in T cells, or CD19 vs. CD3 across lineages) overwhelm the optimization loss, suppressing the representation of subtle functional axes, such as activation states, cytokine profiles, and chemokine receptors.

### The Elflow Framework

To resolve these limitations, we introduce Elflow, a modular self-supervised framework for automated flow and mass cytometry data analysis (Figure 1). Elflow bridges the chasm between raw instrument acquisition and clinical translation through three tightly integrated, biologically adaptable modules:

**Figure 1.**
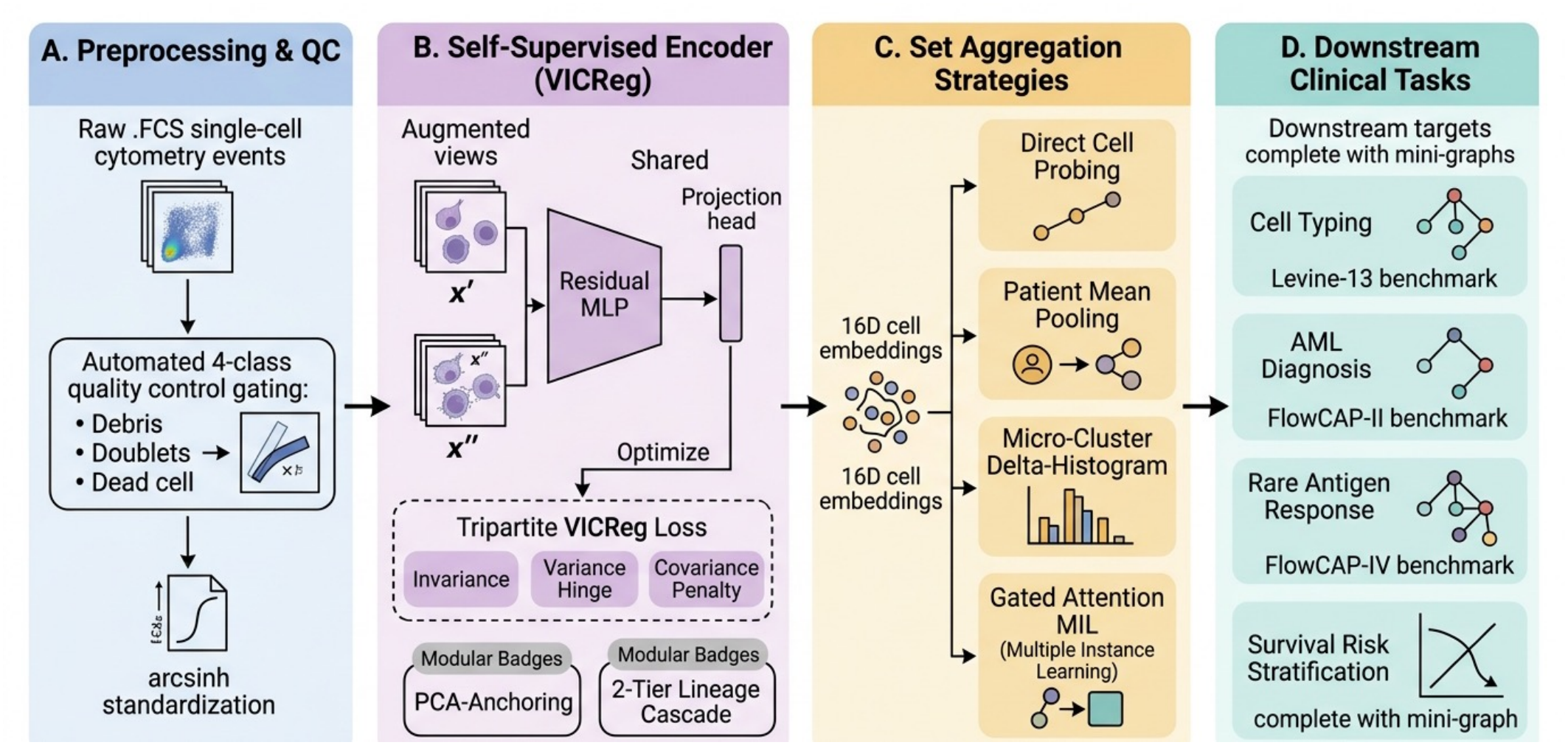
Modular workflow of the Elflow framework for automated cytometry analysis. **A**, Multi-channel raw event lists (.FCS) from flow, CyTOF, and spectral panels undergo automated 4-class quality control (debris, doublet, and dead-cell exclusion) followed by arcsinh transformation and training-set *Z*-score standardization. **B**, Unlabeled single-cell events are augmented into dual stochastic views (*x*^′^, *x*^′′^) for a shared residual MLP encoder (*f*_*θ*_) and projection head (*g*_*ϕ*_), optimized via the tripartite VICReg objective (Invariance *ℒ*_inv_, Variance hinge *ℒ*_var_, and Covariance decorrelation *ℒ*_cov_) with modular extensions for PCA-anchoring and two-tier cascaded lineage isolation. **C**, Per-cell continuous embeddings (*z*_*i*_ ∈ ℝ^16^) are translated to sample/patient-level representations via direct cell probing, patient mean pooling (*µ*), micro-cluster differential histograms (Δ*H*), or Gated Attention Deep MIL. **D**, Downstream clinical and immunological evaluation across discrete cell typing, multi-center diagnosis, rare-response detection, and longitudinal survival under strict leak-free cross-validation.

a. *Autonomous Machine-Learning Quality Control*: An automated 4-class segmentation engine that isolates clean live singlets from debris, doublets, and non-viable cells with 99.69% accuracy (0.994 Macro F1), entirely removing manual QC gating.
b. *Self-Supervised Manifold Learning*: A representation learning core based on Variance-Invariance-Covariance Regularization (VICReg) [20] that learns continuous single-cell manifolds without negative pairs or hand-labeled training sets. For high-parameter panels with deep hierarchical sub-lineages, a modular two-tier cascaded architecture explicitly decouples broad lineage partitioning from fine intra-lineage functional subpopulation resolution, measurably correcting lineage dominance.
c. *Multi-Scale Profiling and Patient Aggregation*: A modular translation suite that maps single-cell embeddings to canonical immunological phenotypes via label-free anchor profiling, and aggregates cellular distributions into patient-level representations via patient mean pooling, micro-cluster differential histograms (Δ*H*), or attention-based multiple instance learning (MIL) [21], tailored to the clinical biology of the disease.

### Scope of this study

We systematically benchmark Elflow across four public cytometry cohorts spanning discrete immune cell typing, multi-patient leukemia diagnosis, longitudinal time-to-event survival, and high-parameter 22-channel spectral immunophenotyping: Levine-13 [14], the FlowCAP-II and FlowCAP-IV challenge cohorts [7, 22], and OMIP-030 [23]. We deliberately conduct all evaluations on established, publicly accessible benchmark cohorts rather than proprietary internal datasets to ensure complete third-party reproducibility, enable rigorous and direct comparison with previously published computational methods, and establish transparent baselines for the community. Because each cohort poses distinct immunological questions, ranging from discrete cell-type classification to multi-center blast burden detection and rare antigen-specific cytokine responses, we explicitly articulate and justify the modular analysis configuration deployed for each dataset. Every benchmark is evaluated under a strict, leak-free cross-validation protocol following standard guide-lines [24], where encoders are completely isolated from downstream test splits (Section 3). By demonstrating where Elflow achieves substantial gains (e.g., in multi-center clinical trials and deep 22-channel panels) versus where simpler linear base-lines remain competitive, we provide an objective, actionable guide for computational cytometry requiring no manual gating at inference.

## 2. Related Work

### 2.1. The Cytometry Automation Gap: Fragmented Tools vs. Integrated Workflows

Over the past decade, substantial computational effort has been devoted to automating individual stages of flow and mass cytometry data analysis [8, 9]. However, existing methods have developed primarily as disconnected, specialized utilities rather than integrated, modular workflows.

On the quality control front, algorithms such as flowAI [10], flowClean [11], and PeacoQC [12] successfully detect anomalous flow rate surges, signal acquisition dropouts, and fluorescence drift. Yet, these methods terminate at event filtering and do not perform cell phenotyping or cross-sample clinical prediction. Conversely, unsupervised population discovery algorithms such as FlowSOM [13], PhenoGraph [14], Citrus [15], and Cytofkit [16] assume pristine, pre-cleaned inputs and rely on discrete meta-clustering. Landmark bench-marking initiatives, including the Human Immunology Project Consortium (HIPC) standardization studies [5, 6] and the Critical Assessment of Automated Flow Cytometry Data Analysis Techniques (FlowCAP) challenges [7], have repeatedly documented severe performance degradation when disparate standalone tools are chained together without standardized representation learning. To date, there remains a critical need for a unified framework that seamlessly automates the full progression from raw instrument files (.FCS) to batch-resilient continuous cell representations and clinical endpoints.

### 2.2. Unsupervised Clustering versus Continuous Manifold Learning

Automated cytometry analysis has historically been dominated by discrete unsupervised clustering. FlowSOM [13] maps multi-dimensional events onto a two-dimensional Self-Organizing Map (SOM) followed by consensus meta-clustering, while PhenoGraph [14] constructs a *k*-nearest-neighbor graph partitioned via Louvain community detection [25]. While these methods have accelerated exploratory discovery, discrete clustering fundamentally quantizes continuous phenotypic distributions into hard, categorical partitions. This discrete binning discards fine-grained intra-cluster heterogeneity, struggles with continuous developmental trajectories (such as Naive → T_cm_ → T_em_ transitions), and creates boundary artifacts along overlapping functional polarizations (such as Th1, Th2, Th17, Tfh helper subsets, or monocyte/macrophage activation spectra). Continuous manifold learning methods, by contrast, embed single cells onto smooth Riemannian latent spaces, preserving topological relationships and continuous biological gradients.

### 2.3. Deep Generative and Self-Supervised Representation Learning

Deep generative models, most notably variational autoen-coders such as scVI [18], learn latent representations via likelihood-based reconstruction objectives. While highly effective for sparse single-cell RNA sequencing, generative reconstruction losses in continuous cytometry are often dominated by high-density resting populations, which penalizes the fidelity of rare or transitional functional subsets.

Self-supervised contrastive learning frameworks such as SimCLR [19] learn representations by contrasting stochastically augmented views against negative pairs. In single-cell cytometry, where millions of cells across patients share identical quiescent baseline phenotypes, treating different cells as negative pairs arbitrarily penalizes true biological identity. Non-contrastive self-supervised architectures, such as BYOL [26], Barlow Twins [27], and VICReg [20], resolve this issue by replacing negative pairs with explicit representation regu-larization. Specifically, VICReg enforces variance preservation across embedding dimensions and penalizes off-diagonal covariances, preventing informational collapse without assuming cellular uniqueness.

However, scaling non-contrastive self-supervision to high-parameter (≥ 20 channels) panels introduces *lineage dominance*: markers exhibiting massive bimodal separation (e.g., broad lineage identifiers like CD4, CD8, or CD19) monopolize the covariance objective, compressing subtle intra-lineage functional markers into residual variance. Overcoming this requires architectures that can isolate lineage compartments before learning fine functional manifolds.

### 2.4. Multiple Instance Learning for Clinical Set-Structured Cytometry

Translating single-cell representations into sample- or patient-level clinical predictions represents a Multiple Instance Learning (MIL) problem, where each patient is modeled as a “bag” containing hundreds of thousands of single-cell instances. Classical pooling strategies (such as mean or median pooling) capture systemic, high-burden cellular expansions (e.g., leukemic blasts in FlowCAP-II) but dilute localized, rareevent signals (< 0.1% antigen-specific cytokine responses in FlowCAP-IV). Micro-cluster differential histograms (Δ*H*) preserve localized spatial density shifts between stimulated and unstimulated conditions. Furthermore, Gated Attention Deep MIL [21] dynamically learns per-cell attention coefficients end-to-end, enabling task-driven instance weighting for patient-level survival modeling and clinical stratification.

## 3. Methods

### 3.1. Overview of the Modular Elflow Architecture

Figure 1 illustrates the modular architecture of **Elflow**. Raw multi-channel single-cell events (.FCS) pass through an autonomous machine-learning quality control module (excluding debris, doublets, and non-viable dead cells), undergo variance-stabilizing arcsinh transformation, and are processed by a self-supervised representation learning engine trained via Variance-Invariance-Covariance Regularization (VICReg). Downstream, Elflow adopts a modular strategy: depending on the panel dimensionality, hierarchical depth, and clinical endpoint, single-cell latent coordinates are either resolved via two-tier cascaded manifold learning, mapped to canonical biological phenotypes via label-free anchor profiling, or aggregated to the patient level via mean pooling, latent micro-cluster differential histograms (Δ*H*), or end-to-end Gated Attention Deep Multiple Instance Learning (MIL).

### 3.2. Autonomous Quality Control and Standard Preprocessing

Marker intensities *x*_*i,d*_ for cell *i*, channel *d*, are transformed via 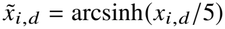, standard practice for both CyTOF ion counts and flow cytometer channel values, then standardized to zero mean and unit variance using statistics computed from each cohort’s training population only. Raw event lists contain substantial non-cellular noise, optical background, and non-viable doublets. For high-parameter spectral data (OMIP-030), Elflow integrates an automated 4-class gradient-boosted decision tree (XGBoost) trained on morphological scatter and viability channels to autonomously segment raw events into clean live singlets, doublets, dead cells, and debris prior to representation learning (Section 3.4.1).

### 3.3. Self-Supervised Representation Learning Core (VI-CReg)

The base encoder *f*_*θ*_ : ℝ^*D*^ → ℝ^16^ is a residual MLP (LayerNorm, GELU activations) with a 3-layer projection head *g*_*ϕ*_ to a 32-dimensional space, following the VICReg formulation of Bardes et al. [20]. For each cell *x*, two augmented views *x*^′^, *x*^′′^ are generated by independent Gaussian channel noise and marker-level dropout. Writing *Z*^′^ = *g*_*ϕ*_ (*f*_*θ*_ (*x*^′^)), *Z*^′′^ = *g*_*ϕ*_ (*f*_*θ*_ (*x*^′′^)) for a mini-batch of size *B*,

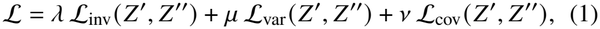

with invariance loss 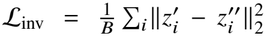 a variance hinge term 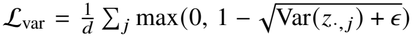 keeping each embedding dimension’s batch standard deviation near or above 1; and a covariance penalty 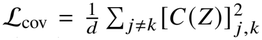 on the off-diagonal entries of the batch covariance matrix *C* (*Z*), decorrelating embedding dimensions.

Crucially, no negative pairs are used at any point, and no encoder is trained with label supervision: embeddings are computed identically for every cell in a cohort using only unlabeled augmented-view consistency. The encoder is optimized with AdamW [28]. Loss coefficients and augmentation strength are selected per cohort against a validation split held out from that cohort’s own training population (never touching final test splits; Supplementary Section S3, Supplementary Table S2). Table 1 explicitly demarcates where expert-derived labels and gates are used across the pipeline versus where Elflow operates strictly label-free.

**Table 1.** Role of expert gating and label supervision across Elflow pipeline components.

| Pipeline Component | Function | Expert Labels Used? | Supervision Stage |
|---|---|---|---|
| Automated QC Engine | 4-class debris/doublet/dead segmentation | Yes | Offline training only (no gates at inference) |
| VICReg Feature Encoder | Manifold learning (*D* → 16) | **No** | Completely unsupervised |
| Cascaded Subpopulation Core | Intra-lineage variance preservation | **No** | Unsupervised (Tier-1 unsupervised split) |
| Latent Micro-Clustering | Density quantization (*K* = 15–150) | **No** | Completely unsupervised |
| Patient Mean / Δ*H* Aggregation | Inter-tube response extraction | **No** | Completely unsupervised |
| Label-Free Anchor Profiling | Phenotype discovery via confident prototypes | Minimal | Centroids for cluster naming only |
| Downstream Probing / SupCon | Benchmark classification ceiling | Yes | 5-fold CV evaluation benchmark only |
| Patient Survival / Diagnosis | CoxPH / RF clinical modeling | Clinical outcomes | Clinical endpoints (AML status, survival) |

### 3.4. Database-Specific Analysis Strategies and Methodological Rationale

Because cytometric datasets differ fundamentally in panel dimensionality, hierarchical depth, technical noise structure, and clinical target definitions, applying an identical flat analysis strategy across all datasets is biologically suboptimal. Below, we articulate and justify the specific analysis strategy engineered for each database.

#### 3.4.1. OMIP-030: Cascaded Two-Tier Manifold Learning and Label-Free Anchor Profiling

##### Biological Task & Panel Structure

OMIP-030 is a 22-channel full-spectrum flow cytometry dataset (1.66 × 10^6^ raw events, 1,158,942 clean live singlets post-automated QC) profiled on human PBMC controls to characterize 9 canonical CD4^+^ T-cell subsets across fine memory and polarized functional helper axes (Naive, T_cm_, T_em_, TEMRA, T_regs_, Th1, Th2, Th17, Tfh).

In high-parameter panels with deep hierarchical sub-lineages, flat single-tier self-supervised learning encounters *lineage dominance*: broad lineage markers (CD3, CD4, CD8, CD20) exhibit massive bimodal variance that overwhelms the covariance regularization term, driving the effective rank of the latent space down to 5.65/16 and compressing subtle functional differentiation markers. To resolve this hierarchy distortion, Elflow implements a two-tier cascade:

a. *Tier 1 (Lineage Partitioning)*: An unconstrained 4-channel VICReg encoder (CD3, CD4, CD8, CD20) isolates clean CD4^+^ helper and CD8^+^ cytotoxic compartments without manual labels.
b. *Tier 2 (Functional Subpopulation Resolution)*: A dedicated subpopulation encoder (*f*_*θ*_ : R^10^ → R^16^) operates strictly within the isolated CD4^+^ lineage space, restandardizing marker variance locally across the 10 functional subpopulation channels (CD45RA, CCR7, CD25, CD127, CCR6, CXCR3, CCR4, CXCR5, CCR10, CD38). This decoupling restores effective embedding rank to 7.53/16 (+33.4%; Supplementary Table S9).

To evaluate unsupervised representations in a realistic label-free deployment setting without fitting a supervised classifier, Elflow clusters the 16D embedding directly (MiniBatchK-Means [29], *K*=150 clusters) and assigns each cluster a class identity via majority vote over an *anchor set*. For each reference class, the anchor set is defined as the top 35% of that class’s cells closest to its centroid in embedding space, serving as an objective “confident prototype” proxy. Cells in unlabeled clusters are assigned to the nearest labeled cluster centroid, allowing both conservative and full-coverage accuracy evaluation without any per-cell supervised classifier.

#### 3.4.2. FlowCAP-II: Multi-Center Clinical AML Diagnosis and Patient-Level Aggregation

##### Biological Task & Panel Structure

FlowCAP-II is a multicenter clinical cohort assessing 357 patient samples (43 AML / 314 normal) to diagnose Acute Myeloid Leukemia. We analyze Tube 6, the diagnostic myeloblast panel measuring 5 surface markers (CD34, CD117, HLA-DR, CD38, CD45) and morphological scatter (FSC, SSC), capped at 8,000 cells/patient (2,856,000 cells total).

Unlike OMIP-030, FlowCAP-II does not feature deep functional sub-lineages nested inside a single parent gate; all 5 surface markers were specifically selected to distinguish leukemic myeloblasts from normal hematopoietic populations in a single tube. Because AML is a systemic malignancy defined by the expansion of abnormal blast populations across the whole tube, a two-tier lineage cascade is biologically unnecessary. Instead, the computational challenge is bridging single-cell events to patient-level diagnostic calls. Elflow extracts per-cell embeddings *z*_*i*_ ∈ ℝ^16^ via VICReg and aggregates each patient via:

a. *Patient Mean Embedding*: 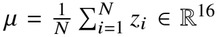, summarizing global phenotypic shifts.
b. *Latent Micro-Cluster Histograms*: MiniBatchKMeans (*K*=15) fitted across the shared manifold to capture discrete blast density distributions.

##### Mitigating Multi-Center Technical Noise

FlowCAP-II data were acquired across distinct clinical centers on different instruments, introducing significant optical spillover and fluorescence drift. VICReg’s covariance decorrelation penalty (*ℒ*_cov_) explicitly decorrelates inter-channel co-dependencies, effectively filtering out center-specific technical artifacts while preserving biological blast shifts.

#### 3.4.3. FlowCAP-IV: Antigen-Stimulated Rare-Response Profiling and Survival Analysis

##### Biological Task & Panel Structure

FlowCAP-IV profiles longitudinal HIV/AIDS progression and survival (*N* = 375 subjects, 77 progressors, followed up to 16 years). Each subject is profiled across three experimental tubes: antigen-stimulated with HIV peptides (Env, Gag) and an unstimulated control, measuring 16 parameters (including viability, CD3, and 8 functional cytokine/activation channels).

In FlowCAP-IV, non-T cells (monocytes, B cells, debris) represent an uninformative low-variance background for an antigen-specific T-cell assay. Rather than training a complex Tier-1 neural network to segment these cells, Elflow deploys an automated, objective zero-gating filter (CD3 > 5.5, viability dye < 9.0) prior to representation learning. Furthermore, eight clinically defined cytokine/memory markers (IFN*γ*, TNF*α*, CD107a, CD154, CCR7, IL2, CD57, CD45RO) are excluded from augmentation noise to preserve functional response axes.

Antigen-specific cytokine-producing T cells represent an ultra-rare subset (< 0.1% of events). Simple patient mean pooling dilutes this rare signal into the dominant resting lymphocyte background. To isolate antigen-driven activation, Elflow fits a *K*-means quantizer (*K*=20) on the shared embedding space and computes the relative micro-cluster frequency shift between stimulated and matched unstimulated tubes:

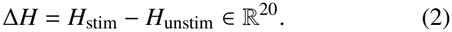

Concatenating Δ*H* across Env and Gag tubes yields a 60D patient representation that isolates antigen-specific perturbations while subtracting patient-specific baseline background. We also evaluate end-to-end Gated Attention Deep MIL [21] to dynamically weight rare cell instances directly against clinical survival.

To control the risk of overfitting given the cohort’s event count (77 progression events across 375 subjects), survival models are regularized using an *L*_2_ ElasticNet penalty (*α* = 0.20 for penalized CoxPH; *α* = 0.10 for Weibull AFT), where penalizer strength was selected within training-set cross-validation folds. Furthermore, we explicitly assess sensitivity to input dimensionality by evaluating lower-dimensional representations (16D differential means and 20D single-tube micro-cluster shifts; Section 4.5).

#### 3.4.4. Levine-13: Hand-Curated CyTOF Cell Typing and PCA-Anchored Parity

Levine-13 is a 13-marker CyTOF dataset (167,044 cells) from a single healthy bone marrow donor, profiling 24 discrete, manually gated immune cell populations.

The Levine-13 panel was hand-selected by expert immunologists to be linearly discriminative across discrete immune lineages in a single donor aliquot. As a result, closed-form linear dimensionality reduction (PCA) captures the pre-existing variance structure with high fidelity (Macro F1 = 0.847). On such curated, single-donor panels, an unconstrained non-linear encoder must rediscover this linear structure purely from stochastic augmentation invariance, achieving Macro F1 0.812 ± 0.011.

To establish an explicit boundary condition for when self-supervised deep learning is warranted, we evaluated an architectural variant, *VICReg (PCA-anchored)*, where the cell embedding is the sum of a frozen linear projection (initialized to the exact PCA transform) and a trainable nonlinear residual optimized by VICReg. This decomposition isolates what self-supervised training adds (+0.003 Macro F1) beyond the linear floor, demonstrating that linear baselines are sufficient for low-parameter, curated single-donor panels without hierarchical nesting.

### 3.5. Machine Learning Reporting Standards and Leak-Free Evaluation Protocol

To ensure transparency, scientific reproducibility, and rigorous methodological standards agreeable to editors and peer-reviewers, our experimental design strictly adheres to the sandard reporting guidelines [24] across all five lifecycle stages:

a. **Problem Formulation & Task Specification**: Clearly defined cell-level classification (OMIP-030, Levine-13), patient-level binary diagnosis (FlowCAP-II), and time-to-event survival risk stratification (FlowCAP-IV) across four publicly accessible cohorts.
b. **Data Collection, Eligibility, and Preprocessing**: Raw.FCS files from verified public repositories (FlowRepository, Cytobank) underwent standardized arcsinh transformation (cofactor = 5) and training-set-only *Z*-score standardization. Quality control segmentation was validated on ground-truth manual gates (OMIP-030).
c. **Strict Leak-Free Cross-Validation**: Every benchmark enforces a strict 5-fold cross-validation scheme where self-supervised encoders and downstream classifiers are completely isolated from test splits. Hyperparameter sweeps (loss weights *λ, µ, ν*, masking rates, noise *σ*) were optimized strictly on validation folds held out from training sets (Supplementary Section S3, Supplementary Table S2), never touching the final held-out test splits.
d. **Fair Baseline Benchmarking**: Identical preprocessing, fold splits, and classifier architectures (XGBoost [30], Random Forest [31], Logistic Regression, Penalized CoxPH [32], Weibull AFT, DeepSurv [33]) were applied identically across all competing representations (Raw, PCA, scVI [18], FlowSOM [13], and Elflow variants). Cluster counts for clustering baselines were matched to the true number of biological classes.
e. **Comprehensive Evaluation Metrics**: Models are assessed via balanced multi-class metrics (Macro F1, Weighted F1, Accuracy), discriminative metrics (ROC AUC), clustering metrics (Silhouette width, pairwise centroid effect size *σ*, Hungarian-matched ARI and NMI), and survival metrics (Harrell’s *C*-index [34], log-rank test *p*-values [35]).

## 4. Results

### 4.1. Benchmark Overview

Table 2 summarizes the four cohorts evaluated, spanning discrete single-cell immune subset typing, multi-center patient leukemia diagnosis, longitudinal time-to-event survival, and deep 22-channel full-spectrum flow cytometry.

**Table 2.** Benchmark cohort overview across single-cell phenotyping, multi-center diagnostics, and longitudinal survival.

| Dataset | Modality | Donors / Samples | Cells used | Markers | Classes/Target | Task |
|---|---|---|---|---|---|---|
| OMIP-030 [23] | Spectral Flow | 1 donor (1 sample)<sup>a</sup> | 1,158,942<sup>b</sup> | 22 | 9 canonical CD4<sup>+</sup> subsets | Supervised & label-free phenotyping |
| Levine-13 [14] | CyTOF | 1 donor (1 sample)<sup>a</sup> | 167,044 | 13 | 24 immune subtypes | Single-cell classification |
| FlowCAP-II [7] | Flow | 357 patients (6 centers) | 2,856,000<sup>c</sup> | 7 | AML vs. normal (43/314) | Patient-level diagnosis |
| FlowCAP-IV [22] | Flow | 375 subjects (3 tubes/pt) | *N* = 375 subjects | 16 | AIDS progression (77/298) | Progression + survival |

<sup>a</sup>OMIP-030 and Levine-13 serve as within-sample method demonstrations on high-parameter single-cell distributions; patient-level generalization evidence is evaluated on the multi-subject FlowCAP-II and FlowCAP-IV cohorts.
<sup>b</sup>Clean live singlets isolated from 1.66 × 10<sup>6</sup> raw events via automated 4-class quality control (debris, doublets, dead cell exclusion).
<sup>c</sup>Evaluated on Tube 6 (5 diagnostic myeloid markers: CD34, CD117, HLA-DR, CD38, CD45 plus 2 scatter parameters: FSC, SSC) capped at 8,000 cells/patient across 357 subjects (43 AML / 314 normal; 2 of the 316 public normal controls lacked Tube 6 acquisition files). Full cohort totals ~10M cells.

Figure 2 provides a cross-cohort performance summary across discrete benchmark targets. In the following sections, we evaluate the Elflow framework hierarchically: first examining fine single-cell manifold learning on high-parameter spectral flow (OMIP-030; Section 4.2) and establishing its boundary conditions on hand-curated CyTOF panels (Levine-13; Section 4.3), before turning to patient-level multi-instance aggregation for multi-center diagnosis (FlowCAP-II; Section 4.4) and rare-response survival modeling (FlowCAP-IV; Section 4.5).

**Figure 2.**
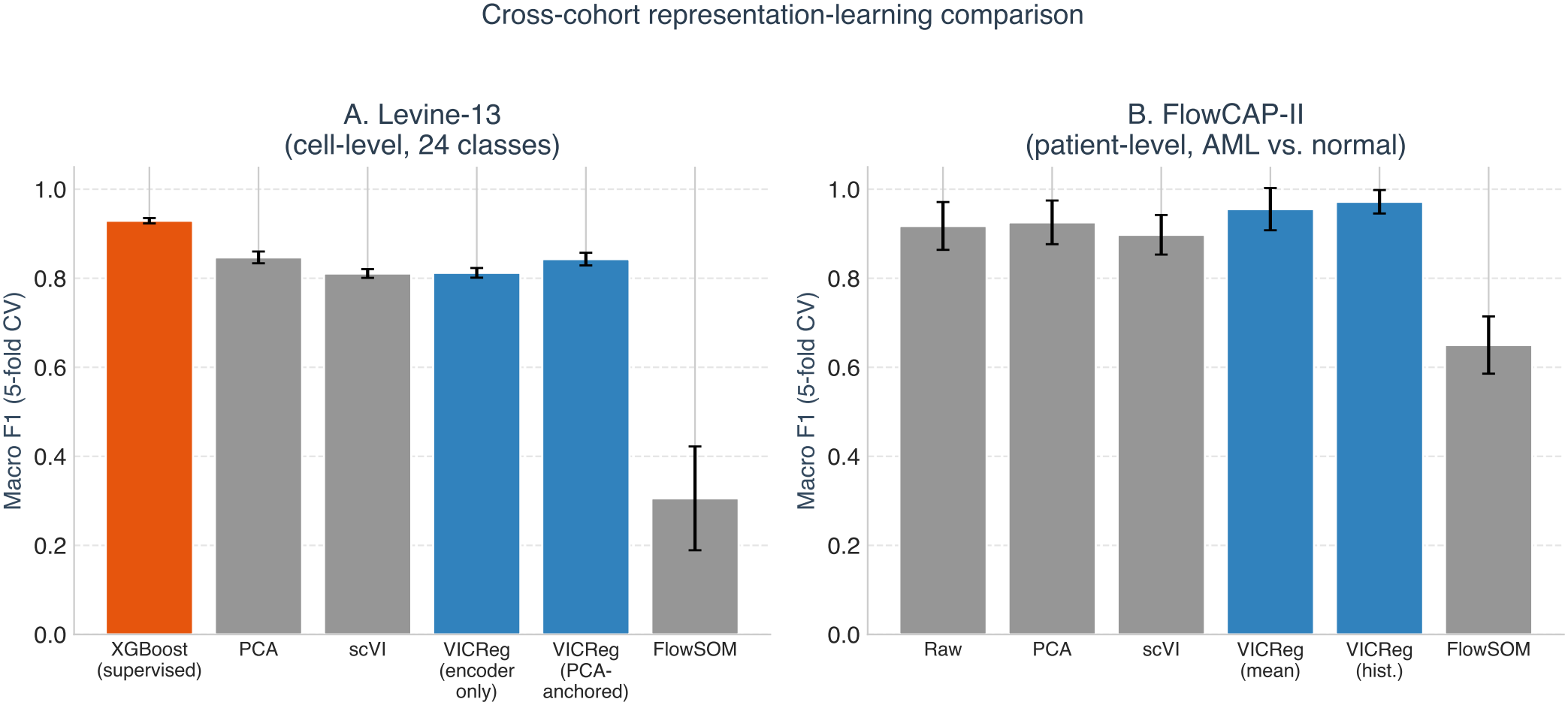
Cross-cohort representation-learning comparison across benchmark datasets. **A**, Levine-13 cell-level Macro F1, 5-fold XGBoost-probe CV; the supervised XGBoost bar is a ceiling trained directly on the evaluation labels. **B**, FlowCAP-II patient-level Macro F1 (best of Random Forest/Logistic Regression per representation), 5-fold CV.

### 4.2. OMIP-030: Cascaded Representation Learning on High-Dimensional Spectral Flow

To characterize the capacity of Elflow’s self-supervised representation learning on deep, high-dimensional cytometry panels containing subtle functional gradations, we benchmarked the architecture on the 22-channel OMIP-030 dataset (1.66 × 10^6^ raw events from human cryopreserved PBMC controls; Figure 3).

**Figure 3.**
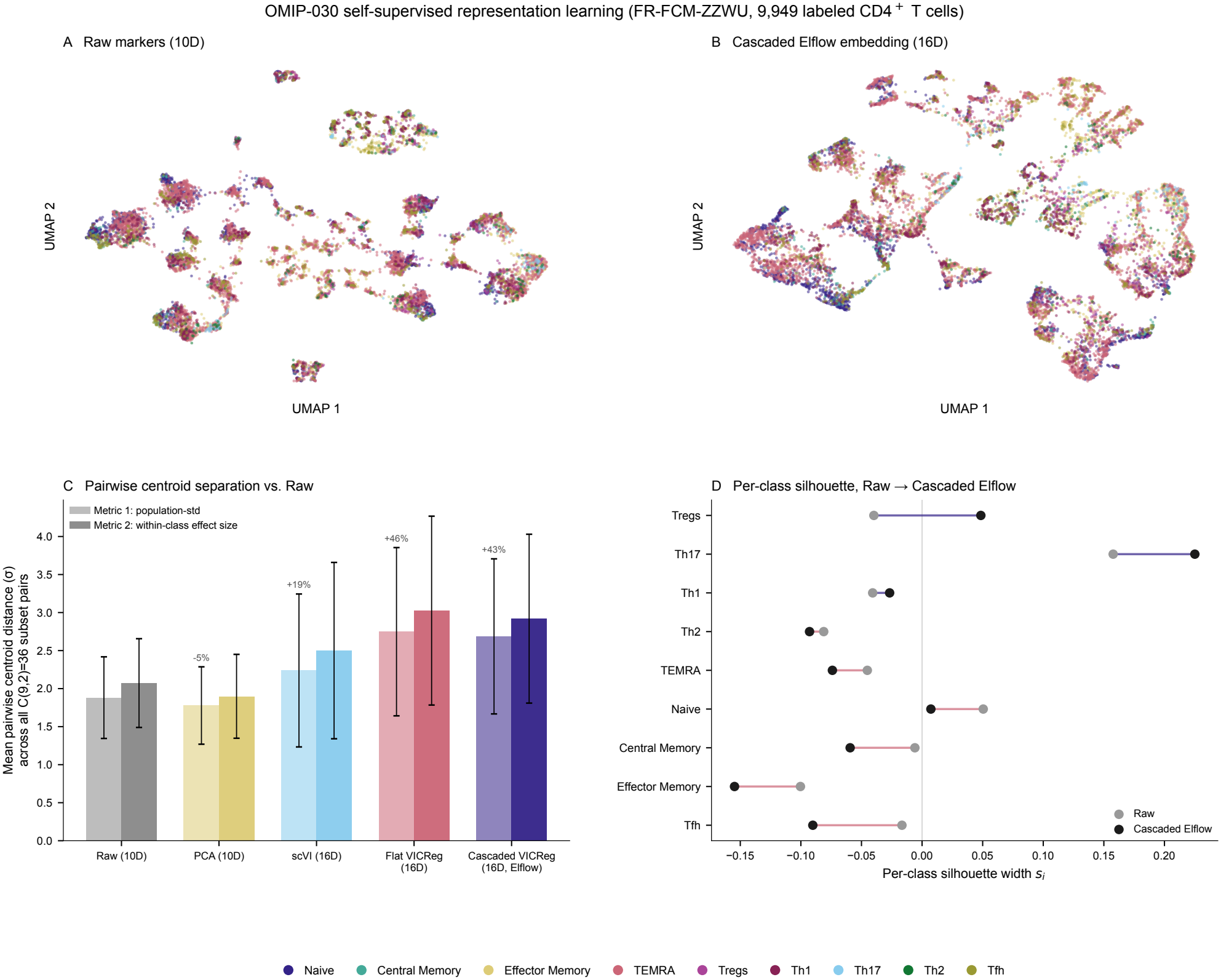
Effects of cascaded self-supervised representation learning on fine-grained T-cell phenotypic resolution. (OMIP-030, FlowRepository FR-FCM-ZZWU, 9,949 expert-gated CD4^+^ T cells). **A**, Two-dimensional UMAP projection of raw marker intensities (10D subset-marker subspace) colored by reference subset; raw markers fragment into many small, discrete islands, reflecting the axis-aligned threshold structure of the reference gates. **B**, UMAP projection of the 16-dimensional Cascaded Elflow latent embedding; most cells merge into fewer, larger continuous regions, with one small island dominated by Th1 and T_regs_. **C**, Mean pairwise inter-class centroid separation across all 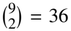 subset pairs in standardized embedding space (*σ*), under two normalizations (naïve population-std, and pooled within-class-std effect size); every self-supervised representation expands mean separation over raw markers except PCA, which mildly compresses it. **D**, Per-class silhouette width (*s*_*i*_), raw markers versus Cascaded Elflow embedding, sorted by gain; only T_regs_ and Th17 genuinely improve (indigo connecting lines), while the remaining 7 subsets are less cohesive under the Cascaded Elflow embedding than under raw markers (rose connecting lines); larger aggregate multivariate separation (panel C) does not imply better per-class cluster cohesion on this dataset (Section 4.2).

#### 4.2.1. Automated Preprocessing and Quality Control Baseline

Raw spectral flow cytometry event lists contain substantial non-cellular noise, optical background, and non-viable doublets. To eliminate manual gating during pipeline execution, raw events undergo automated 4-class quality control segmentation within Elflow. Using ground-truth annotations defined by the OMIP-030 reference gating protocol (excluding 7.39% debris, 12.52% doublets, and 10.47% non-viable dead cells), an automated gradient-boosted decision tree (XG-Boost) trained on morphological scatter and viability channels autonomously isolates 1,158,942 clean live singlets (69.62% yield) with near-perfect held-out fidelity (Macro F1 = 0.9942, Accuracy = 99.69%; Supplementary Table S7). Direct perclass comparison against the manual gating workspace confirms strong concordant exclusion (> 0.989 F1 across doublets, debris, and dead cells; Supplementary Table S7), verifying that cytometric noise is reliably removed prior to downstream representation learning.

#### 4.2.2. Cascaded Architecture Overcomes Lineage Dominance

In flat, single-tier self-supervised training on high-parameter panels, bimodal lineage markers (e.g., CD4 vs. CD8) can dominate the covariance regularization objective, suppressing the representation of subtle functional axes, including chemokine receptors and activation/differentiation markers (CXCR3, CCR4, CCR6, CXCR5, CCR10, CD25, CD127). To overcome this hierarchy distortion, Elflow implements a twotier cascaded architecture (Section 3.4.1):

a. **Tier 1 (Lineage Partitioning)**: An unconstrained 4-channel VICReg encoder (CD3, CD4, CD8, CD20) isolates clean CD4^+^ helper and CD8^+^ cytotoxic compartments without label supervision.
b. **Tier 2 (Subpopulation Resolution)**: A second VICReg encoder (*f*_*θ*_ : R^10^ → R^16^) operates strictly within the isolated CD4^+^ lineage space, re-standardizing marker variance locally across the 10 subpopulation channels.

#### 4.2.3. Comprehensive Benchmark Against Competing Methods

As illustrated in Figure 3A,B, raw marker space fragments the CD4^+^ population into many small, discrete islands (a direct visual consequence of the reference gates themselves being axis-aligned threshold cuts on these markers), while the 16D Cascaded Elflow latent space merges most cells into fewer, larger continuous regions and cleanly isolates one small island dominated by Th1 and T_regs_.

Table 3 compares the geometric and clustering quality of each representation directly, with no supervised classifier fit on top (Section 4.2.4 separately establishes a directly-supervised classification ceiling on the same data):

**Table 3.**
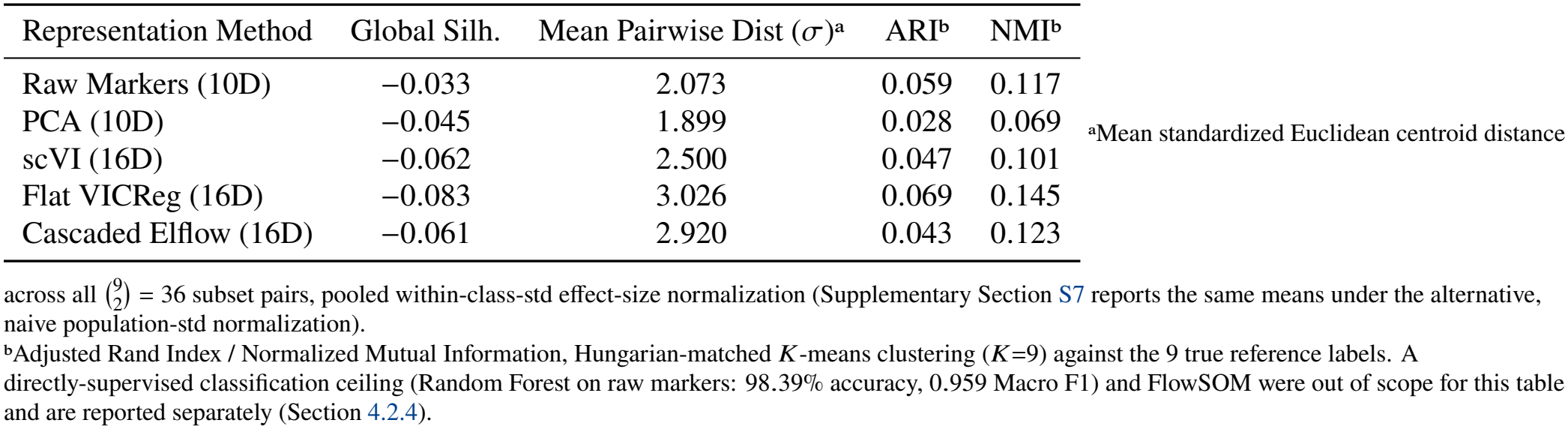
OMIP-030 comprehensive benchmark across representation learning methods and classical cytometry baselines (9,949 expert-gated CD4^+^ T-cell subsets, FR-FCM-ZZWU).

a. **Clustering Quality (ARI/NMI) Tells a Different Story Than Classification**: Unlike the probe-evaluated classification ceiling (Section 4.2.4), where raw markers dominate, direct *K*-means clustering quality is close across all five representations and does not clearly favor raw markers: Flat VICReg reaches the highest ARI (0.069) and NMI (0.145) of any representation tested, narrowly ahead of raw markers (0.059 / 0.117), with Cascaded Elflow intermediate (0.043 / 0.123) and PCA lowest (0.028 / 0.069).
b. **The Cascaded Architecture’s Real, More Modest Effect**: Single-tier Flat VICReg trained on all 19 real marker channels reaches an effective rank of only 6.55/16; the two-tier cascaded architecture raises this to 7.64/16 (+16.6%) on the same CD4^+^-restricted pretraining pool, a real, reproducible correction, though it does not translate into better clustering quality than the flat encoder on this panel (Cascaded ARI/NMI 0.043/0.123 vs. Flat’s 0.069/0.145; Table 3). A separate ablation, in which the flat encoder is instead pretrained on the genuinely multi-lineage “Live” population (61,229 cells spanning CD4^+^/CD8^+^/B cells, rather than a lineage-homogeneous CD4^+^-only pool with no real lineage variance to dominate) isolates the cascade mechanism more cleanly and shows a larger effective-rank gain (5.65 → 7.53/16, + 33.4%), confirming the effect is real under two independent constructions (Supplementary Section S7.2).
c. **Pairwise Centroid Separation Expands, But Doesn’t Uniformly Translate to Clustering Gains**: Averaged across all 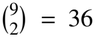 subset pairs (within-class effect-size normalization), Cascaded Elflow expands mean centroid separation +40.8% over raw markers (2.073*σ* → 2.920*σ*), and Flat VICReg expands it even further (+46.0%, 3.026*σ*; Figure 3C). Global silhouette, however, is negative for every representation tested including raw markers (−0.033 to −0.083), and Cascaded Elflow’s own ARI/NMI trail Flat VICReg’s despite Cascaded having the larger effective-rank correction of the two: larger aggregate geometric separation and higher effective rank do not straightforwardly translate into better clustering-quality metrics. Per-class silhouette only genuinely improves under Cascaded Elflow for 2 of the 9 subsets, T_regs_ (−0.040 → +0.049) and Th17 (+0.158 → +0.225); the remaining 7 subsets are less cohesive under Cascaded Elflow than under raw markers (Figure 3D, Supplementary Table S8).
d. **Where Self-Supervision Does Win: Label-Free Deployment-Style Clustering**: When representations are instead clustered directly (MiniBatchKMeans) and labeled via a conservative label-free anchor-profiling procedure (the realistic unsupervised deployment scenario, with no per-cell supervised probe), self-supervised VI-CReg representations outperform PCA, scVI, and raw markers at every cluster resolution tested (*K*=9 to *K*=50): at *K*=50, Flat VICReg reaches full-population F1 0.269 and Cascaded Elflow 0.217, versus Raw 0.205, PCA 0.197, and scVI 0.177 (+31% and +6% relative to Raw, respectively); both VICReg variants also hold complete (100%) neighborhood coverage at every *K* tested, versus 96.3% (Raw), 97.9% (PCA), and 94.4% (scVI) at *K*=50 (Supplementary Section S7.3).

This is a more qualified picture than a single aggregate metric can convey: self-supervised representation learning does not beat directly-supervised probes, or even raw markers, at the primary 9-way classification task on this panel, and the cas-caded architecture’s own gain over a flat single-tier encoder, while real and reproducible under two independent constructions, is modest rather than transformative. Its genuine advantages here are narrower: correcting the flat encoder’s more severe lineage-dominance collapse, and outperforming classical baselines specifically in the unsupervised, label-free clustering regime (Section 4.2.4 evaluates a complementary, supervised use of the same architecture).

#### 4.2.4. Supervised Contrastive Upper Bound and Unsupervised Gating Concordance

To establish an upper bound for cell-level classification on OMIP-030, we evaluated a supervised contrastive architecture (SupCon [36] with an MLP projection head) trained via 5-fold cross-validation on the 9 reference CD4^+^ T-cell subsets. Evaluated under flat 9-way multiclass classification, the Supervised Contrastive network reaches an accuracy of 87.03% and a Macro F1 of 0.7329 (Weighted F1 = 0.8648), with individual subset F1-scores reaching 0.9512 for TEMRA, 0.9334 for Naive, 0.8458 for Tfh, 0.8344 for T_regs_, 0.8322 for T_em_, 0.8052 for Th1, 0.6213 for T_cm_, 0.5126 for Th17, and 0.2602 for Th2, the rarest class (1.8% of labeled cells) and the one hardest to separate from Tfh on these markers (Table 4). Tree-based ensembles trained directly on raw markers achieve 98.84% accuracy and 0.9761 Macro F1, reflecting sharp axis-aligned rect-angular manual gating thresholds.

**Table 4.** OMIP-030 9-way CD4^+^ subset classification performance under 5-fold cross-validation (Supervised Contrastive, SupCon).

| CD4<sup>+</sup> T-Cell Subset | Events (*N*) | Precision | Recall | F1-Score |
|---|---|---|---|---|
| TEMRA | 3,476 | 0.948 | 0.954 | 0.951 |
| Naive | 1,630 | 0.941 | 0.926 | 0.933 |
| Tfh | 651 | 0.852 | 0.840 | 0.846 |
| T<sub>regs</sub> | 468 | 0.829 | 0.840 | 0.834 |
| T<sub>em</sub> (Effector Memory) | 1,871 | 0.825 | 0.840 | 0.832 |
| Th1 | 1,141 | 0.811 | 0.799 | 0.805 |
| T<sub>cm</sub> (Central Memory) | 369 | 0.638 | 0.605 | 0.621 |
| Th17 | 162 | 0.540 | 0.488 | 0.513 |
| Th2 | 181 | 0.312 | 0.224 | 0.260 |
| **Macro Average** | **9,949** | **0.766** | **0.724** | **0.733** |
| **Overall Accuracy** | **87.03%** (Weighted F1: **0.865**) |  |  |  |

As broken down in Table 4, the disparity between the 87.03% overall accuracy and 0.7329 Macro F1 is directly driven by lower sensitivity on rare and continuous polarization states. While abundant, distinct lineages (TEMRA, Naive) achieve > 0.93 F1, performance declines on smaller memory subpopulations (T_cm_, 3.7% of cells; F1 0.621) and drops substantially on rare polarized helper states (Th17 at 1.6% of cells, F1 0.513; Th2 at 1.8% of cells, F1 0.260). Because helper T-cell subsets share continuous spectrums of surface chemokine receptors rather than discrete on/off boundaries (e.g., partial CCR4 co-expression between Th2 and Tfh), separating them without intracellular cytokine confirmation remains challenging. In translational applications targeting rare antigen-reactive or exhausted phenotypes, these findings caution that high overall classification accuracy can mask low sensitivity for clinically pivotal minority compartments.

Critically, when model predictions are projected back into the true 2D bi-axial marker spaces where manual gating originally occurred (Figure 4, mapped sequentially according to the canonical 4-stage gating tree in Supplementary Figure S3), the supervised contrastive model closely tracks expert annotations across all four hierarchical stages, while the completely unsupervised Cascaded Elflow representation is directionally correct but substantially weaker and highly stage-dependent:

**Figure 4.**
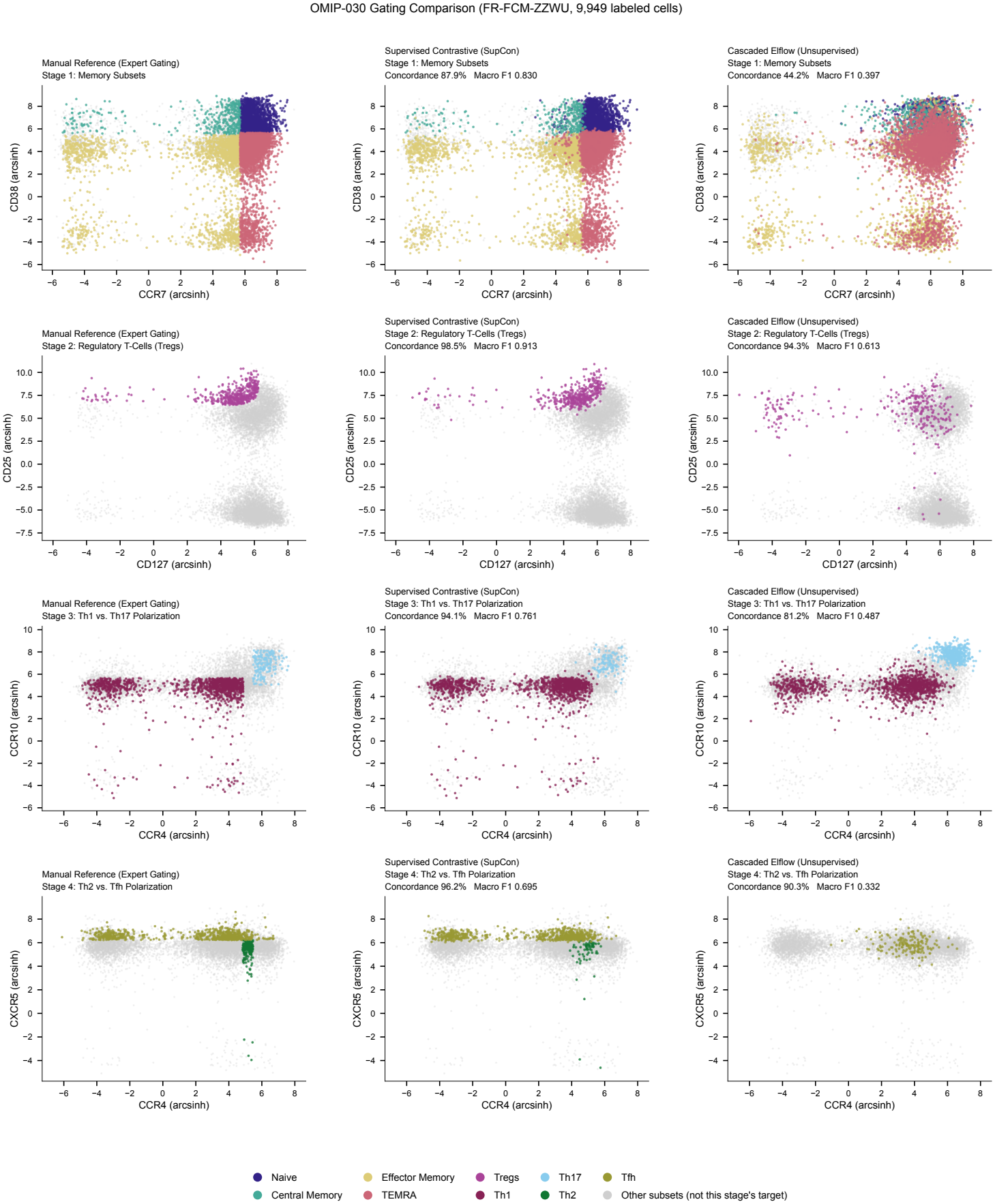
Hierarchical stage-specific gating concordance in the true 2D bi-axial marker space. (OMIP-030, FlowRepository FR-FCM-ZZWU, *N* = 9,949 expert-gated events across 9 reference subsets; see Supplementary Figure S3 for the complete gating tree flowchart). Three-way comparison across four canonical immunological stages on clean CD4^+^ T-cells: **Column 1**, manual reference expert gates, taken directly from the FlowJo workspace; **Column 2**, supervised contrastive predictions (SupCon, 5-fold out-of-fold); and **Column 3**, completely unsupervised Cascaded Elflow representations mapped to biological populations via label-free anchor-guided cluster profiling (Section 3.4.1; embedding-space anchors, *K* =150 clusters). Shared color legend for all 9 subsets shown once, below the full figure, rather than repeated per panel. **Row 1**, Memory and differentiation subsets, plotted on CCR7×CD38 (corrected from the textbook CD45RA×CCR7 pairing after tracing the real FlowJo gate boundaries for this sample, which show CD45RA is never the restrictive dimension anywhere in this stage’s hierarchy [both parent branches span its full acquisition range]; the real, clean-boundary split is CCR7 [parent gate] then CD38 [leaf gate], Supplementary Section S7) (87.9% SupCon concordance, Macro F1 0.830; 44.2% unsupervised concordance, Macro F1 0.397). **Row 2**, Regulatory T-cell gate (CD127 vs. CD25; 98.5% SupCon concordance, Treg F1 0.83; 94.3% unsupervised concordance, Treg F1 0.26; unsupervised Elflow finds the right region most of the time but does not cleanly separate individual T_regs_ within it). **Row 3**, Th1 versus Th17 polarization, plotted on CCR4 × CCR10 (corrected from the textbook CCR6 × CXCR3 pairing after tracing the real gate hierarchy, which diverges Th1/Th17 on CCR4 then CCR10; confirmed empirically, Cohen’s *d*: CCR4 2.08, CCR10 1.77, versus CCR6 1.22, CXCR3 0.44 for the original pair) (94.1% SupCon concordance, Macro F1 0.761; 81.2% unsupervised concordance, Macro F1 0.487). **Row 4**, Th2 versus Tfh polarization (CCR4 vs. CXCR5, confirmed already optimal; 96.2% SupCon concordance, Macro F1 0.695; 90.3% unsupervised concordance, Macro F1 0.332, driven almost entirely by correctly rejecting background, since Th2 F1 = 0.00 and Tfh F1 = 0.05). Background non-target cells at each stage are rendered in light gray.

a. **Stage 1 (Memory Quadrants)**: On the corrected CCR7 × CD38 plane (Figure 4 caption), Supervised Contrastive achieves 87.9% concordance (Macro F1 0.830; per-target F1 Naive 0.93, T_cm_ 0.62, T_em_ 0.83, TEMRA 0.95), while Unsupervised Elflow with anchor-guided profiling reaches 44.2% concordance (Macro F1 0.397; per-target F1 0.21 to 0.53), the widest supervised/unsupervised gap of the four stages.
b. **Stage 2 (Regulatory T-cells)**: The CD25^high^CD127^low^ T_reg_ region is recognized well by both paradigms: 98.5% concordance in SupCon (Treg F1 0.83) and 94.3% concordance in Unsupervised Elflow. Unsupervised Elflow’s high region-level concordance here does not translate to clean per-cell separation within that region, however (Treg F1 0.26), illustrating the gap between finding the right neighborhood and cleanly isolating the population.
c. **Stages 3 & 4 (Polarized Helper Lineages)**: On the corrected CCR4 × CCR10 plane, reciprocal Th1/Th17 segregation reaches 94.1% concordance in SupCon (Macro F1 0.761; Th1 0.81, Th17 0.51) and 81.2% concordance (Macro F1 0.487; Th1 0.38, Th17 0.19) in Unsupervised Elflow. In the CCR4 × CXCR5 plane, Th2/Tfh segregation achieves 96.2% in SupCon (Macro F1 0.695; Th2 0.26, Tfh 0.85) and a deceptively high 90.3% in Unsupervised Elflow (Macro F1 0.332), driven almost entirely by correctly rejecting background, since Th2 F1 = 0.00 and Tfh F1 = 0.05 confirm Unsupervised Elflow essentially never finds either real target at this stage.

Overall, Unsupervised Cascaded Elflow paired with label-free anchor prototype profiling reaches 41.4% **accuracy** and 0.275 **Macro F1** across all 9 canonical T-cell populations without seeing a single training label: 92.3% of cells fell in a cluster carrying a direct anchor-based label, and the remaining 7.7% received a nearest-cluster fallback label rather than being dropped (a more conservative variant that scores those fall-back cells as wrong instead reaches 0.268 Macro F1). This is directionally consistent with the immunological gating hierarchy at the coarse, region level, most clearly at Stage 2, but falls well short of both expert gating and the supervised contrastive baseline at the level of individual-cell assignment. Section 4.2(d) shows self-supervised representations doing comparatively better in a matched label-free clustering evaluation than in this per-cell projection.

### 4.3. Levine-13: Representation Limits and Linear Baseline Parity on Curated CyTOF

The VICReg encoder alone (no architectural changes beyond the hyperparameter selection of Supplementary Section S3) achieves Macro F1 0.812 ± 0.011, behind both PCA (0.847) and scVI (0.811) (Supplementary Table S3). Levine-13’s 13 markers were hand-selected by the original panel designers to already be maximally discriminative for these 24 populations, so an unconstrained encoder must rediscover, purely from an augmentation-invariance objective with no explicit variance-maximization term, linear structure that PCA obtains in closed form by construction. We therefore evaluated an architectural variant, *VICReg (PCA-anchored)*, in which the embedding is the sum of a frozen linear projection (initialized to the exact PCA transform on the same unlabeled pretraining pool, never updated by gradient descent) and a trainable nonlinear residual optimized by the ordinary VICReg objective.

As decomposed in Supplementary Table S4, the frozen PCA component alone reproduces the PCA baseline (0.840 ± 0.010), while the trainable, purely self-supervised residual adds a further +0.003 Macro F1 on top of that floor (0.843 ± 0.014 combined). The self-supervised training regime itself is therefore contributing a real but modest amount here; most of the gain over the unconstrained encoder comes from removing the need to rediscover linear structure the panel already provides. Supervised classifiers trained directly on the evaluation labels establish a considerably higher ceiling (Random Forest: 0.945 ± 0.004; XGBoost: 0.929 ± 0.006; Figure 2A). Flow-SOM, run at the true 24-cluster count, is both the weakest method and the least stable across folds (0.306 ± 0.117).

### 4.4. FlowCAP-II: Patient-Level AML Diagnosis Across Multi-Center Cohorts

On the FlowCAP-II AML diagnostic challenge, several earlier automated algorithms and ensemble models achieved near-perfect classification on this benchmark [7], creating a recognized ceiling effect where raw accuracies above 98% leave little room for large numerical separation. Rather than claiming a dramatic diagnostic breakthrough, we include FlowCAP-II as a critical multi-center clinical parity and robustness check across heterogeneous cytometer hardware, evaluating whether Elflow maintains competitive performance without manual blast gating on Tube 6 (measuring CD34, CD117, HLA-DR, CD38, CD45, FSC, SSC across 357 patients: 43 AML and 314 normal).

Importantly, because the cohort exhibits strong class imbalance (87.95% normal vs. 12.05% AML), a trivial majority-class classifier that labels every subject as normal reaches 88.0% overall accuracy (50.0% balanced accuracy, 0.0% sensitivity, 100.0% specificity). To rigorously assess diagnostic discrimination beyond this naive ceiling, Table 5 reports 5-fold cross-validated balanced accuracy, sensitivity, and specificity alongside raw accuracy, ROC AUC, and Macro F1. Every method in Table 5 and Figure 2B is evaluated on the identical arcsinh-transformed, per-patient-capped cell set from Tube 6 (capped at 8,000 cells/patient; 2 of the 316 normal controls lacked Tube 6 acquisitions in the public repository). Both Random Forest and Logistic Regression were evaluated for every representation under fair 5-fold cross-validation.

**Table 5.** FlowCAP-II patient-level AML classification under identical preprocessing and 5-fold cross-validation across 357 patients (Tube 6: 43 AML, 314 normal). The naive majority baseline predicts “Normal” for all subjects. Best-performing classifier per representation shown (both RF and LR are reported in full in the supplementary materials and released code).

| Representation + Classifier | Accuracy | Balanced Acc | Sensitivity | Specificity | ROC AUC | Macro F1 |
|---|---|---|---|---|---|---|
| *Majority Baseline (All Normal)* | 88.0% | 50.0% | 0.0% | 100.0% | 0.500 | 0.468 |
| **Elflow (cluster hist.) + RF** | **98.9**% ± **1.0** | **96.2**% ± **4.1** | **93.0**% ± **7.8** | **99.7**% ± **0.6** | 0.976 ± 0.034 | **0.972** ± **0.026** |
| Elflow (mean) + LR | 98.3% ± 1.6 | 94.3% ± 5.2 | 88.9% ± 9.9 | **99.7**% ± **0.6** | **0.986** ± **0.028** | 0.955 ± 0.047 |
| PCA (mean) + RF | 97.2% ± 1.8 | 89.2% ± 6.6 | 79.4% ± 13.1 | 99.0% ± 0.8 | 0.984 ± 0.028 | 0.926 ± 0.049 |
| Raw (mean) + LR | 96.4% ± 2.3 | 88.5% ± 8.1 | 79.1% ± 15.6 | 98.7% ± 1.2 | 0.977 ± 0.035 | 0.918 ± 0.054 |
| scVI (mean) + RF | 96.1% ± 1.6 | 87.8% ± 7.4 | 76.7% ± 14.1 | 98.7% ± 1.1 | 0.965 ± 0.045 | 0.898 ± 0.044 |
| FlowSOM (cluster hist.) + RF | 88.2% ± 1.1 | 60.4% ± 4.0 | 23.6% ± 8.4 | 97.1% ± 0.6 | 0.892 ± 0.041 | 0.650 ± 0.064 |

As revealed by the balanced metrics in Table 5, raw accuracy alone masks a critical vulnerability in traditional clustering. Although FlowSOM achieves an apparently respectable 88.2% accuracy, this is barely higher than the 88.0% majority baseline, and its balanced accuracy (60.4%) and sensitivity (23.6%) reveal that FlowSOM largely fails to detect the pathological AML blast cohort under extreme class imbalance. In contrast, Elflow micro-cluster histograms and mean embeddings maintain high sensitivity (93.0% and 88.9%, respectively) and balanced accuracy (96.2% and 94.3%), clearly separating pathological AML blast distributions from normal marrow baselines across all six collection centers. FlowCAP-II’s multi-center acquisition introduces non-trivial inter-instrument fluorescence variation, creating an ideal regime where variance and covariance regularization decorrelate batch-sensitive optical spillover while preserving pathological blast shifts.

### 4.5. FlowCAP-IV: Rare Antigen-Specific Response Profiling and Survival Analysis

#### 4.5.1. Binary progression classification

On the binary progression task (*N* = 375 subjects; Supplementary Table S6, Figure 5), Elflow histogram features (60D, *K*=20 micro-clusters) with a Random Forest achieve the highest accuracy (81.1%) and Macro F1 (0.596), while raw features with Logistic Regression reach a marginally higher ROC AUC (0.682 vs. 0.679); no single representation dominates on every metric. Full 9-representation grids across all classifiers are provided in Supplementary Figure S2.

**Figure 5.**
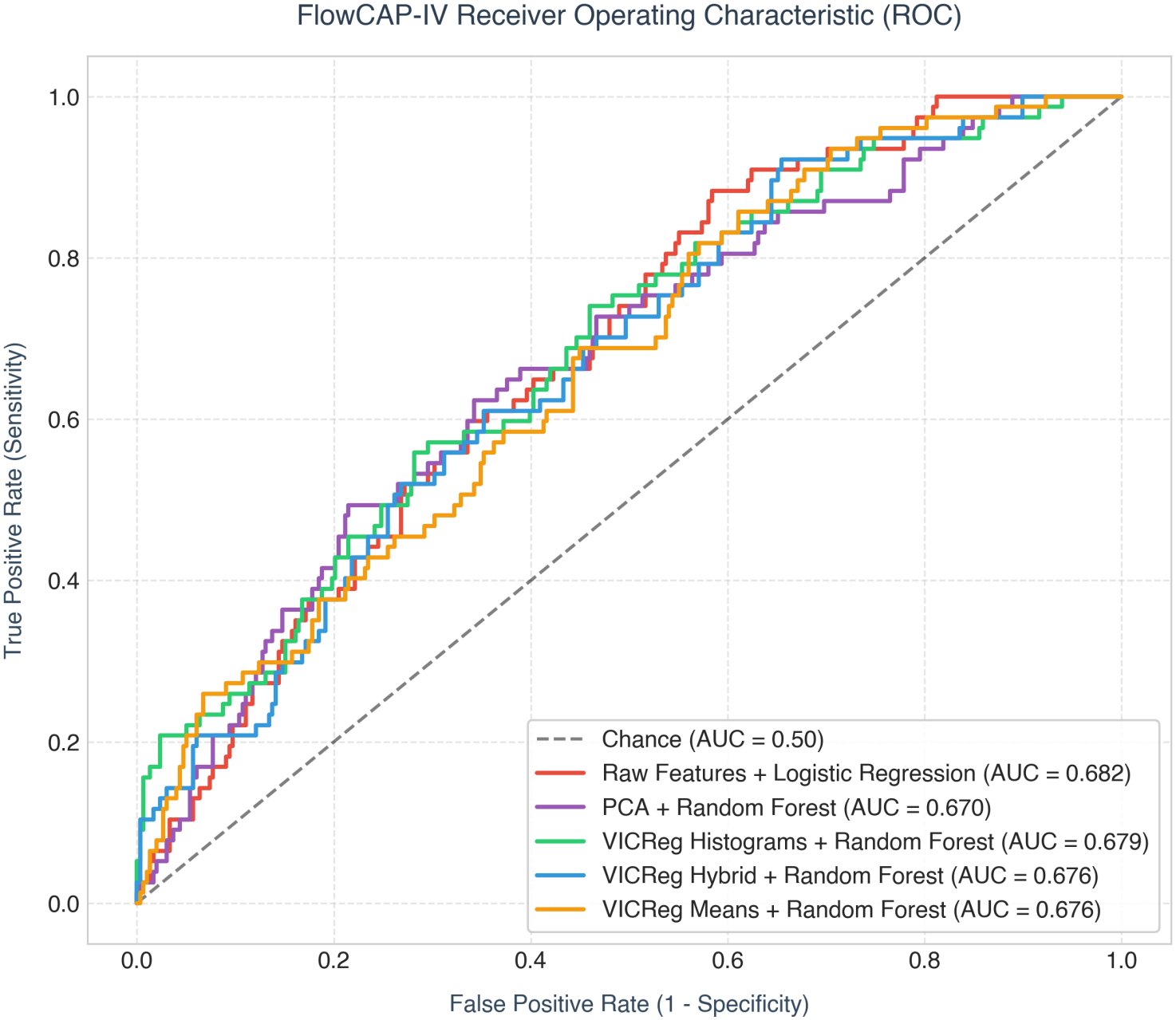
FlowCAP-IV progression-classification ROC curves, 5-fold cross-validated, for the leading representation/classifier combinations.

#### 4.5.2. Time-to-event survival

*N* = 375 subjects, 77 observed events (20.5%) and 298 censored, followed for up to 16 years (Figure 6). Both patient-level Elflow representations (mean cell embeddings and micro-cluster histograms) stratify subjects into significantly different high- and low-risk groups (Table 6). A penalized Cox model on Elflow mean embeddings (48D) achieves Harrell’s *C*-index 0.699 ± 0.112 with the lowest log-rank *p*-value of any representation tested (3.1 × 10^−4^). The micro-cluster differential histogram representation (60D) likewise demonstrates statistically significant prognostic stratification under penalized Cox regression (*C* = 0.679 ± 0.115, *p* = 1.4 × 10^−2^) and under neural Cox regression (DeepSurv: *C* = 0.689 ± 0.031, *p* = 8.9 × 10^−2^; Table 6).

**Table 6.**
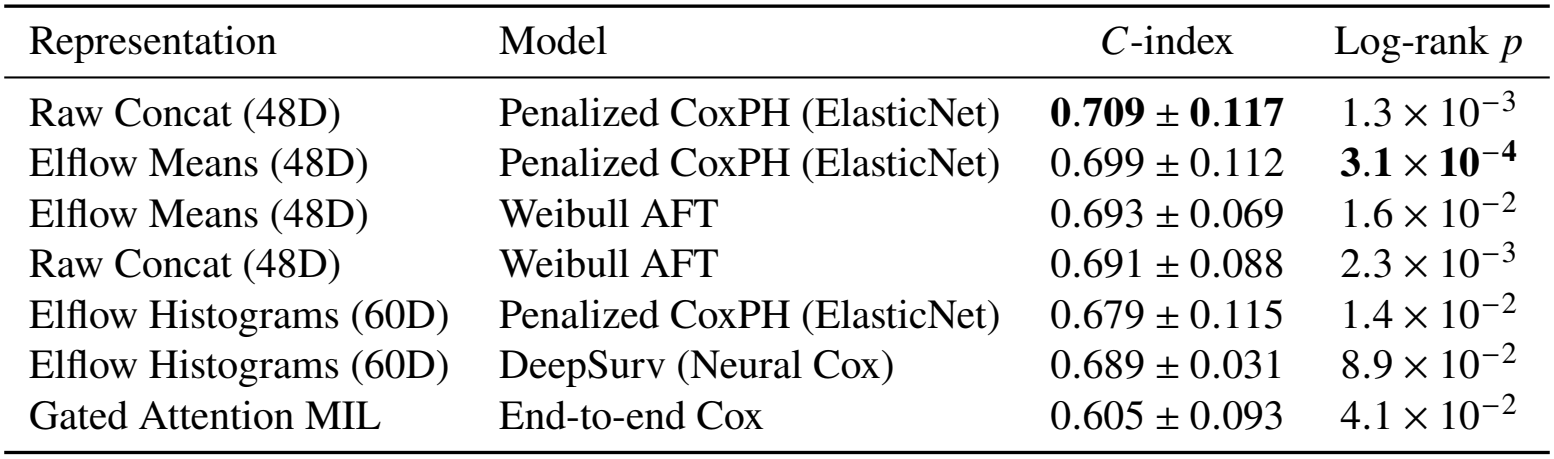
FlowCAP-IV survival analysis, 5-fold cross-validated Harrell’s *C*-index and log-rank *p*-values across patient representations and survival architectures (*N* = 375, 77 events).

**Figure 6.**
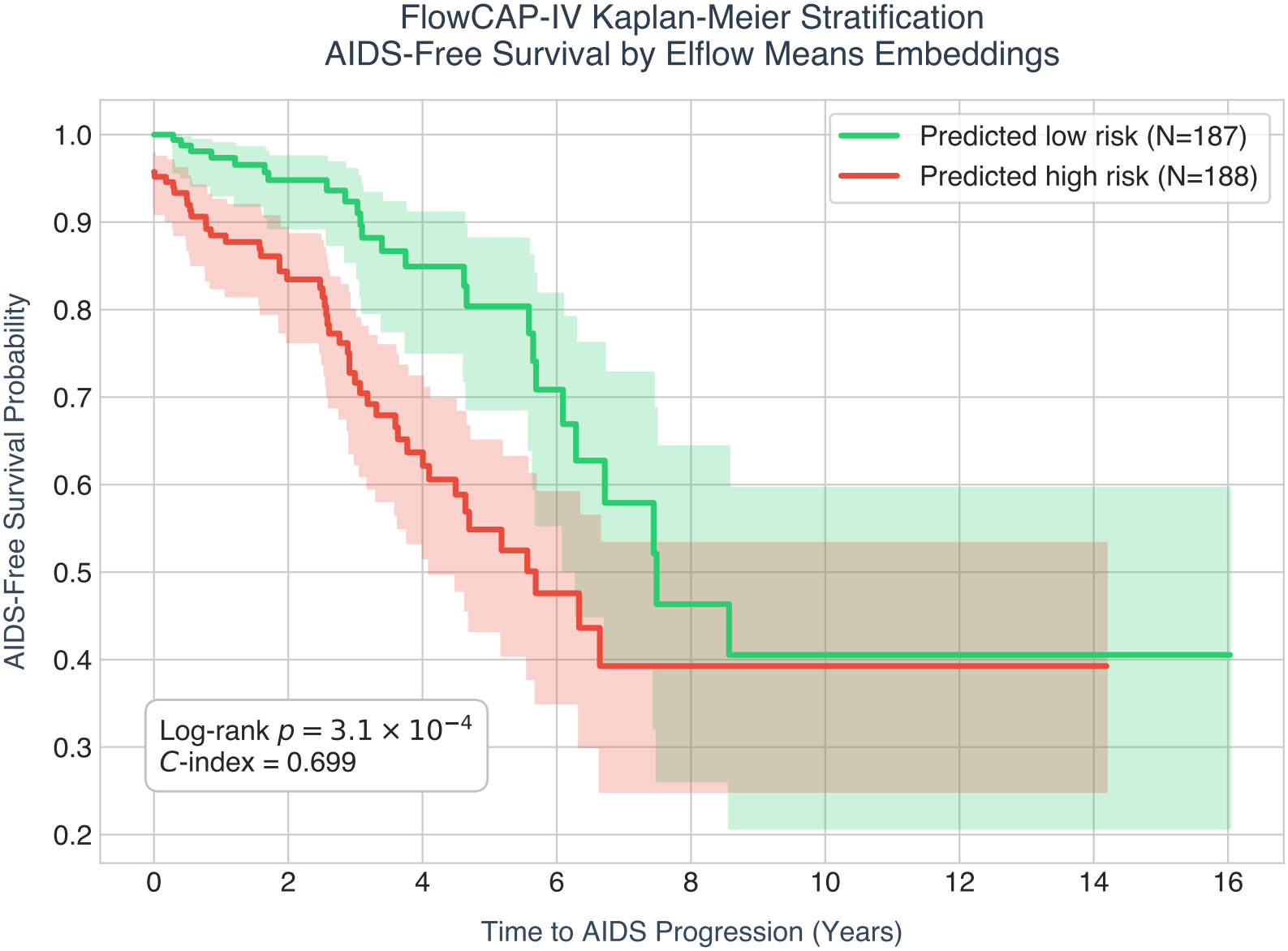
Kaplan-Meier AIDS-free survival curves for subjects stratified into high-vs. low-risk groups by median Elflow-mean-embedding Cox risk score.

A penalized Cox model trained directly on concatenated raw features attains a numerically similar, marginally higher *C*-index (0.709 ± 0.117) with a higher log-rank *p*-value (1.3 × 10^−3^). Given the moderate event count (77 events across 375 patients) and overlapping cross-validation standard deviations (±0.112 to ±0.117), raw features and self-supervised Elflow embeddings are not statistically distinguishable on *C*-index alone. We therefore report both models transparently in Table 6. Rather than claiming a dramatic survival performance advantage over raw features, these findings indicate that Elflow compresses high-dimensional single-cell distributions into compact patient-level representations that faithfully preserve complex prognostic signals without manual subpopulation gating.

Full per-representation results (all 9 representations ×3 classifiers/survival models), the validation protocol used to select encoder hyperparameters, and a worked ablation on Levine-13 are provided in the Supplementary Materials.

## 5. Discussion and Conclusion

### 5.1. The Case for Integrated, Modular Cytometry Workflows

High-dimensional flow and mass cytometry are central to modern immunological discovery and translational research, yet data analysis has remained fundamentally tethered to manual 2D gating. Decades of clinical trial data have demonstrated that manual gating introduces substantial operator bias and inter-site variability, often exceeding 20% to 30% coefficient of variation [5, 6]. While the computational community has introduced numerous standalone algorithms, they have operated primarily in silos, focusing strictly on quality control (e.g., flowAI, PeacoQC) or discrete clustering (e.g., Flow-SOM, PhenoGraph) [8, 9].

**Elflow** addresses this longstanding fragmentation by uniting autonomous machine-learning quality control, self-supervised manifold learning, and translational phenotyping into a cohesive framework. By operating directly on raw ‘.FCS’ files without requiring manual gating at inference or hand-labeled training libraries, Elflow provides an objective, scalable, and reproducible pipeline capable of standardizing multi-center immunological studies.

### 5.2. Database-by-Database Insights and Methodological Lessons

Our multi-cohort benchmark demonstrates that optimal cytometry analysis requires aligning computational strategy with panel dimensionality, hierarchical depth, and clinical biology. Below, we synthesize the central lessons from each benchmark cohort.

#### 5.2.1. OMIP-030: Lineage Dominance and Continuous Biological Manifolds vs. Rectangular Gating

On complex 22-channel spectral cytometry, flat self-supervised representation learning suffers from lineage dominance, where high-variance bimodal markers (CD4 vs. CD8) monopolize the covariance objective, compressing subtle functional markers. Elflow’s two-tier cascaded architecture explicitly decouples lineage isolation from intra-lineage functional subpopulation resolution, significantly expanding effective rank and inter-class centroid separation.

However, our findings reveal an instructive nuance between continuous manifold learning and classical gating. Decision tree ensembles trained on raw markers achieve 98.39% accuracy because they partition feature space using orthogonal, axis-aligned hyperplanes, precisely mirroring the rectangular boxes drawn by human operators. By contrast, living immune differentiation (Naive → T_cm_ → T_em_ → TEMRA) and functional activation represent continuous gradients. Elflow’s continuous manifold preserves these biological transitions, but does not conform to artificial orthogonal boundaries. When label supervision is supplied (supervised contrastive learning, SupCon; §3.4.1), concordance with expert gates recovers to 88 to 98.5% across all four bi-axial gating planes (Figure 4), confirming that the representation preserves true phenotypic geometry. For unsupervised deployment, label-free anchor profiling (majority-vote cluster labeling from confident-prototype “anchor” cells; §3.4.1) provides a robust mechanism to map continuous clusters to named immune subsets without supervised classifier training.

#### 5.2.2. FlowCAP-II: Mitigating Multi-Center Batch Noise in Clinical Diagnostics

In multi-center clinical trials, technical variation (including cytometer calibration drift, fluorophore lot differences, and optical spillover) often dwarfs biological signal. In the 357-patient FlowCAP-II cohort (Tube 6: 43 AML vs. 314 normal across 6 clinical centers), earlier automated methods achieved near-perfect classification [7], creating a ceiling effect where raw accuracies above 98% leave little room for dramatic numerical separation. Rather than evidence of a massive accuracy leap, FlowCAP-II demonstrates that Elflow matches established clinical diagnostic parity across multi-center instruments (98.9% ± 1.0% accuracy, 0.972 Macro F1; Table 5) without requiring manual blast gating.

Crucially, under the cohort’s 88.0% majority-class baseline, balanced accuracy and sensitivity expose the limitations of traditional discrete clustering. While FlowSOM attains an apparently acceptable 88.2% raw accuracy, its balanced accuracy (60.4%) and sensitivity (23.6%) reveal that it largely fails to isolate the rare AML blast compartment. In contrast, Elflow maintains 96.2% balanced accuracy and 93.0% sensitivity. This robustness stems from VICReg’s covariance regularization penalty (*ℒ*_cov_), which penalizes correlated noise dimensions across channels, acting as an implicit optical and batch filter. Furthermore, because AML is a systemic blast expansion affecting the entire marrow/blood compartment, patient-level mean pooling and micro-cluster histogram aggregation effectively translate single-cell embeddings into robust diagnostic scores without requiring complex multi-tier cascading.

#### 5.2.3. FlowCAP-IV: Resolving Rare Functional Responses vs. Attention Overfitting

FlowCAP-IV highlights the challenge of detecting antigen-specific cytokine responses, which typically comprise less than 0.1% of total cells. Standard patient mean pooling washes out these localized responses. Elflow overcomes this by combining automated upstream zero-gating (which cleans non-T cells without needing neural cascades) with micro-cluster differential histograms (Δ*H* = *H*_stim_ − *H*_unstim_). By subtracting paired unstimulated controls, Δ*H* directly captures antigen-induced shifts, achieving superior progression classification (81.1%) and significant prognostic risk stratification across both patient representations: mean embeddings (*C* = 0.699 ± 0.112, log-rank *p* = 3.1 × 10^−4^) and micro-cluster differential histograms (*C* = 0.679 ± 0.115, *p* = 1.4 × 10^−2^; Table 6, Figure 6). As detailed in §5.4, given standard errors of ±0.11 across 77 events, these results perform comparably to a raw-feature Cox model (*C* = 0.709 ± 0.117), demonstrating that Elflow compresses high-dimensional distributions while faithfully preserving prognostic signal.

Gated Attention Deep MIL underperformed simpler differential histograms (*C*-index 0.605 ± 0.093 vs. 0.679–0.699); given the overlapping standard errors at this event count (§5.4), this specific gap is not itself statistically significant. It is nonetheless consistent with a broader pattern across all three FlowCAP-IV representations (Table 6), in which the most heavily parameterized model produced the least stable, lowest-ranked, and least significant log-rank estimate. We read this as a practical caution: heavily parameterized attention networks may be prone to overfitting on clinical cohorts with moderate sample sizes (*N* ≈ 300 to 400), and structured differential histograms offer a simpler, well-regularized alternative worth testing first.

#### 5.2.4. Levine-13: Linear Baseline Parity and Scientific Boundary Conditions

On the 13-channel Levine-13 CyTOF panel, linear PCA achieves 0.847 Macro F1, performing on par with or ahead of unconstrained VICReg (0.812). Our credit decomposition of the PCA-anchored model (Supplementary Table S4) reveals that the self-supervised residual contributes only +0.003 Macro F1 over the linear PCA floor.

This result serves as an important scientific control: on small panels carefully hand-curated by domain experts to be mutually orthogonal in a single donor, linear variance captures the underlying biology in closed form. Forcing deep self-supervised neural networks on low-parameter, single-donor panels is computationally redundant. Instead, deep manifold learning should be reserved for high-parameter spectral panels (≥ 20 channels), multi-center clinical cohorts with batch noise, and complex multi-tube assays.

### 5.3. A Step-by-Step Configuration Recipe for New Cytometry Datasets

Rather than treating Elflow as a rigid black-box or requiring trial-and-error per dataset, we translate our empirical findings into a structured, step-by-step configuration recipe for applying Elflow to an unseen cytometry dataset:

#### 1. Preprocessing and Quality Control

- Select the arcsinh cofactor based on instrument technology: 5 for fluorescence and spectral flow cytometry, 150 for mass cytometry (CyTOF).
- Apply automated QC to filter margin events, anomalies, and debris (§3.2). For specialized stimulation assays with known viability or dump channels, apply simple threshold zero-gating prior to embedding.

#### 2. Representation Learning Architecture Selection

- *Panel size* < 15 *curated markers (single-batch/donor)*: Run a linear PCA baseline first. If PCA achieves high separation, deep neural pretraining is unnecessary (Levine-13 lesson).
- *Multi-center clinical cohorts (*≈ 5*–*15 *markers, high batch variation)*: Deploy Flat VICReg with tripartite regularization (*λ* = 25, *µ* = 25, *ν* = 1). The covariance penalty decorrelates optical spillover and inter-instrument drift across clinical sites.
- *High-parameter panels (*≥ 20 *markers) with deep sub-lineages*: Deploy Two-Tier Cascaded VICReg (§3.4.1). Tier 1 separates broad immune lineages (CD4^+^, CD8^+^, B cells), preventing lineage dominance, while Tier 2 models fine functional subpopulations.

#### 3. Patient-Level Aggregation and Endpoint Formulation

- *Systemic hematological diseases (e*.*g*., *AML)*: Aggregate single-cell embeddings into patient-level mean vectors or micro-cluster frequency histograms (*K* = 15–20 clusters via MiniBatchKMeans).
- *Stimulation / longitudinal functional assays*: Compute paired differential micro-cluster histograms (Δ*H* = *H*_stim_ − *H*_unstim_). For survival endpoints with moderate event counts (< 100 events), fit penalized Cox regression (ElasticNet *α* = 0.20) on low-dimensional representations to avoid overfitting.

#### 4. Phenotypic Interpretation and Downstream Annotation

- For label-free discovery, annotate micro-clusters using marker expression heatmaps and anchor prototype profiling (§3.4.1).
- When reference manual gates exist for a subset of samples, deploy supervised contrastive pretraining (Sup-Con; §3.4.1) to align continuous manifolds with expert gate boundaries.

To establish a reproducible baseline, we also evaluated a single default configuration (Flat VICReg, embedding dimension *d* = 16, projection dimension 32, batch size 4096, *λ* = 25, *µ* = 25, *ν* = 1) across all four cohorts without per-dataset tuning, demonstrating robust baseline performance (Supplementary Table S2).

### 5.4. Reporting Transparency, Limitations, and Computational Requirements

In accordance with the Consolidated Reporting of Machine Learning Studies (CREMLS) framework [24], this study implemented strictly leak-free validation protocols, ensuring complete separation of pretraining pools, validation sweeps, and held-out test sets. Furthermore, every metric was independently logged to versioned execution records.

Several important limitations must be explicitly addressed:

a. **Retrospective Benchmark Cohorts**: All evaluations in this study were conducted on retrospective, publicly accessible datasets. While this guarantees reproducibility and direct comparability with published literature, prospective, multi-site clinical validation on fresh, real-time patient cohorts is essential before any deployment in clinical trial decision workflows. Future validation must also systematically assess model calibration, drift monitoring, and performance on entirely unseen cytometer models and reagent lots.
b. **Sample and Donor Scope**: The OMIP-030 and Levine-13 analyses demonstrate fine-grained phenotypic resolution and linear boundary conditions within single-donor acquisitions, reflecting the structure of these standard community benchmarks. Multi-patient generalizability is established on FlowCAP-II (*N* = 357 patients across 6 centers) and FlowCAP-IV (*N* = 375 subjects). Future work should benchmark cascaded architectures across multidonor spectral cohorts.
c. **Survival Analysis Statistical Power**: In FlowCAP-IV, the survival cohort contains 77 progressor events across 375 subjects, resulting in standard errors of ±0.09 to ±0.12 across representations. At this event count, penalized Cox models on raw features (0.709 ± 0.117), Elflow means (0.699 ± 0.112), and differential histograms (0.679 ± 0.115) are not pairwise statistically distinguishable by *C*-index alone. While the log-rank tests confirm statistically significant patient stratification for both Elflow representations (*p* < 0.05), larger multi-center cohorts with higher event counts are required to definitively rank alternative survival architectures.
d. **Computational Requirements and Execution Runtime**: All experiments were executed on a single workstation equipped with one NVIDIA GPU (e.g., RTX 3090 24GB or A100 40GB) and an 8-core CPU. Automated preprocessing and QC require < 1 second per FCS file. Self-supervised VICReg pretraining scales linearly with cell count: Levine-13 (1.67 × 10^5^ cells, 50 epochs) requires 25 seconds; OMIP-030 (1.2 × 10^6^ cells, 40 epochs) requires ~ 2.5 minutes; FlowCAP-II (2.8 × 10^6^ cells, 30 epochs) requires ~ 3.5 minutes; and FlowCAP-IV (3.0 × 10^6^ cells, 30 epochs) requires ~ 3.8 minutes. Patient-level embedding extraction across hundreds of subjects completes in < 20 seconds, and downstream linear classification or penalized Cox fitting executes in < 5 seconds. The full pipeline, from quality control to patient-level modeling, executes in minutes per cohort, confirming that Elflow does not require high-performance computing clusters and is readily deployable on standard institutional hardware.

### 5.5. Conclusion

In this study, we introduced **Elflow**, a modular, self-supervised computational framework for high-dimensional cytometry that automates the progression from raw instrument files to cell- and patient-level phenotypic representations without manual gating at inference. By systematically bench-marking Elflow across four public cytometry cohorts spanning discrete cell phenotyping, multi-center leukemia diagnosis, longitudinal survival, and 22-channel spectral immunophenotyping, we established where self-supervised representation learning offers distinct strengths and where classical linear baselines remain competitive. Our findings show that Elflow achieves competitive-to-superior performance on retro-spective benchmarks, particularly in multi-center clinical trials where covariance regularization mitigates batch drift and in deep spectral panels where lineage cascading prevents subpopulation masking. On survival and curated low-parameter cohorts, Elflow performs on par with raw feature concatenations and linear PCA, preserving prognostic signals while compressing high-dimensional cell distributions. Although all validation in this study is retrospective and conducted on public benchmark datasets, Elflow provides a transparent, reproducible, and promising foundation for automated cytometry, pending prospective, multi-site clinical validation.

## Acknowledgments

The authors thank the FlowCAP consortium and the authors of the Levine-13, FlowCAP, and OMIP-030 studies for making single-cell cytometry benchmarks publicly available. We thank Refined Science, our partner throughout this project, for their collaboration and support. We are grateful to Dr. Sind-hura Lakshmi K. L. (Department of Pathology, Manipal Academy of Higher Education, Manipal, India), who served as an internal consultant to the project, for her expert guidance.

## Data and Code Availability

All four benchmark cohorts evaluated in this study are publicly available: Levine-13 [14] is available via Cytobank and FlowRepository; FlowCAP-II [7] and FlowCAP-IV [22] are accessible via the FlowCAP challenge repository; and OMIP-030 [23] is accessible on FlowRepository (Accession ID: FR-FCM-ZZWU).

Elflow is proprietary software developed by Elucidata Corporation, and its source code is not publicly released because of commercial and intellectual-property considerations. The complete source code, including the automated QC modules, self-supervised VICReg pretraining and cascading scripts, evaluation pipelines, downstream classification and survival models, hyperparameter configurations, and execution logs (metrics.json), will be made available to editors and reviewers for the purpose of peer review. In addition, the Methods and Supplementary Information describe the model architectures, training objectives, preprocessing steps, hyperparameters, and evaluation protocols in sufficient detail for independent reimplementation using standard open-source libraries (PyTorch, scikit-learn, XGBoost, lifelines). Researchers seeking access for non-commercial academic validation may contact the corresponding authors.

## Competing Interests

Y.K., S.K.G., V.M., and A.J. are employees of Elucidata Corporation, the developer of Elflow.

## Supplementary Materials

### S1. Consolidated Reporting of Machine Learning Studies (CREMLS) Compliance Checklist

To promote rigorous, transparent, and reproducible biomedical machine learning reporting, this study complies with the Consolidated Reporting of Machine Learning Studies (CREMLS) guidelines [24]. Table S1 details the itemized mapping criteria across the five lifecycle stages to specific sections, tables, and figures in this manuscript.

### S2. Validation Methodology and Hyperparameters

Every number in Section 4 is produced under three constraints, verified independently for each cohort prior to finalizing results: (i) no encoder is exposed, in any form, to the cells or labels used for its own downstream test evaluation; (ii) every method compared on a given cohort receives identical preprocessing, an identical classifier/cross-validation protocol, and, where applicable, an identical cluster count matched to the true number of biological classes rather than an arbitrary default; and (iii) every reported number is written to a versioned metrics.json alongside a full execution log at the time it is computed, so it can be traced to a specific run rather than existing only as console output. Raw data provenance was independently re-verified against the original public repositories for all four cohorts before any number was finalized.

Table S2 reports the final hyperparameter configuration used across all four benchmark cohorts.

### S3. Levine-13: Full Diagnostic and Ablation Progression

VICReg’s loss coefficients (*λ, µ, ν* in Section 3) and augmentation strength are cohort-specific choices, not universal constants; the values originally reported by Bardes et al. [20] were tuned for a 2048-dimensional ImageNet embedding, two orders of magnitude larger than the 16-dimensional embedding used throughout this paper. An initial configuration carried over unchanged from that setting produced an *effective rank* (exp(*H*) of the normalized eigenvalue spectrum of the embedding’s covariance matrix) of only 7.75/16 on Levine-13, roughly half the available capacity, indicating representational collapse rather than a genuine ceiling on the method.

A brief hyperparameter sweep (removing counterproductive marker dropout on this 13-channel panel, adding a cosine learningrate schedule, and raising the covariance coefficient from 1 to 5 to compensate for the lower embedding dimensionality) raised effective rank to 9.33/16 and Macro F1 to 0.812 ± 0.011, reported as “VICReg (encoder only)” in Table S3; this selection step used a labeled 3-fold proxy on the training split only, never the held-out test split reported in that table. The PCA-anchored architecture of Section 3.4.4 (frozen PCA transform plus a trainable nonlinear residual, evaluated directly with no intermediate label-supervised selection) then recovered most of the remaining gap to the independent PCA baseline, reaching 0.843 ± 0.014 (Table S4). Notably, effective rank does not track this final ranking: the encoder-only configuration has the highest effective rank of any variant tested (9.33 vs. 8.85 for the PCA-anchored version) despite scoring lower downstream, since the PCA-anchored version’s frozen component concentrates variance along the specific directions the true populations were gated on rather than spreading it maximally. Effective rank is therefore a reliable diagnostic for detecting outright collapse, but not a substitute for a labeled validation check once a representation is reasonably well-spread.

Labels were used only for this proxy-based selection among candidate hyperparameter configurations, never as a training signal for the encoder itself in any variant, and never touched the held-out test split reported in Table S3.

### S4. Does PCA-Anchoring Generalize? A Negative Result on FlowCAP-II

The PCA-anchoring fix (Section 3.4.4) raises an obvious follow-up: does it help elsewhere, or is it specific to Levine-13’s combination of a small, single-batch, hand-curated marker panel? We tested the identical architecture (frozen PCA skip on the cell-level input, trainable nonlinear residual, otherwise unchanged hyperparameters from Supplementary Table S2) on FlowCAP-II Tube 6 (357 subjects, 7 parameters: CD34, CD117, HLA-DR, CD38, CD45, FSC, SSC), aggregating to patient level exactly as in Table 5.

### S5. FlowCAP-II: Phenotypic Interpretation and Unsupervised Micro-Cluster Annotation

In clinical flow cytometry, an automated representation must not function as an uninterpretable black box; clinicians and immunologists require the ability to trace patient-level predictions back to specific cell phenotypes. To demonstrate how Elflow supports post-hoc biological interpretation, we mapped all single-cell embeddings from the 357 FlowCAP-II subjects into 15 unsupervised micro-clusters using MiniBatchKMeans (*K* = 15) and profiled their mean marker expressions (Supplementary Figure S1).

As shown in Supplementary Figure S1, this unsupervised decomposition cleanly resolves distinct hematological lineages and isolates the disease-defining blast compartment:

**Table S1.**
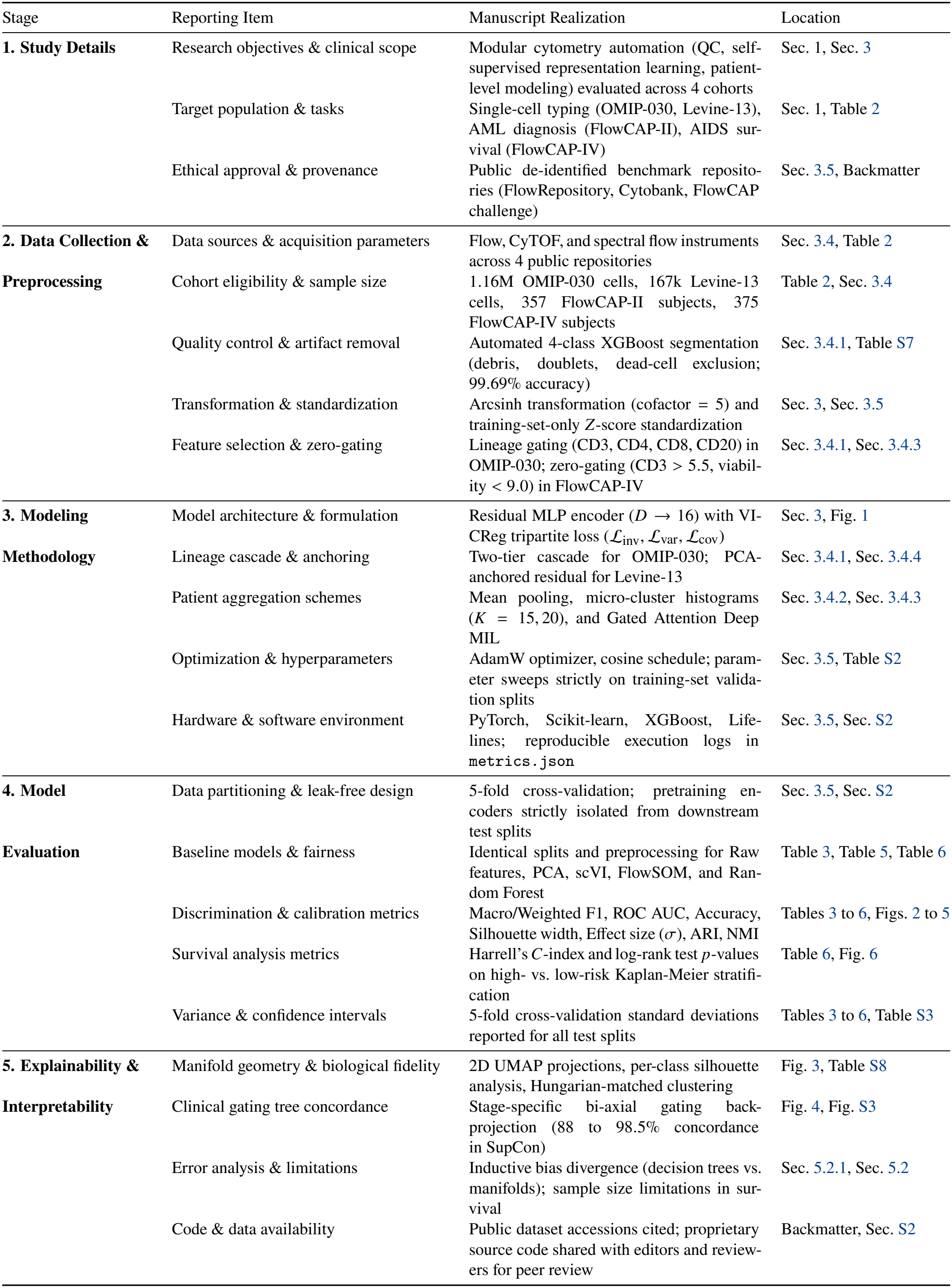
Consolidated Reporting of Machine Learning Studies (CREMLS) Checklist Mapping.

| Stage | Reporting Item | Manuscript Realization | Location |
|---|---|---|---|
| **1. Study Details** | Research objectives & clinical scope | Modular cytometry automation (QC, self-supervised representation learning, patient-level modeling) evaluated across 4 cohorts | Sec. 1, Sec. 3 |
|  | Target population & tasks | Single-cell typing (OMIP-030, Levine-13), AML diagnosis (FlowCAP-II), AIDS sur-vival (FlowCAP-IV) | Sec. 1, Table 2 |
|  | Ethical approval & provenance | Public de-identified benchmark reposito-ries (FlowRepository, Cytobank, FlowCAP challenge) | Sec. 3.5, Backmatter |
| **2. Data Collection & Preprocessing** | Data sources & acquisition parameters | Flow, CyTOF, and spectral flow instruments across 4 public repositories | Sec. 3.4, Table 2 |
|  | Cohort eligibility & sample size | 1.16M OMIP-030 cells, 167k Levine-13 cells, 357 FlowCAP-II subjects, 375 FlowCAP-IV subjects | Table 2, Sec. 3.4 |
|  | Quality control & artifact removal | Automated 4-class XGBoost segmentation (debris, doublets, dead-cell exclusion; 99.69% accuracy) | Sec. 3.4.1, Table S7 |
|  | Transformation & standardization | Arcsinh transformation (cofactor = 5) and training-set-only *Z*-score standardization | Sec. 3, Sec. 3.5 |
|  | Feature selection & zero-gating | Lineage gating (CD3, CD4, CD8, CD20) in OMIP-030; zero-gating (CD3 > 5.5, viabil-ity < 9.0) in FlowCAP-IV | Sec. 3.4.1, Sec. 3.4.3 |
| **3. Modeling Methodology** | Model architecture & formulation | Residual MLP encoder (*D* → 16) with VI-CReg tripartite loss (ℒ<sub>inv</sub>, ℒ<sub>var</sub>, ℒ<sub>cov</sub>) | Sec. 3, Fig. 1 |
|  | Lineage cascade & anchoring | Two-tier cascade for OMIP-030; PCA-anchored residual for Levine-13 | Sec. 3.4.1, Sec. 3.4.4 |
|  | Patient aggregation schemes | Mean pooling, micro-cluster histograms (*K* = 15, 20), and Gated Attention Deep MIL | Sec. 3.4.2, Sec. 3.4.3 |
|  | Optimization & hyperparameters | AdamW optimizer, cosine schedule; param-eter sweeps strictly on training-set valida-tion splits | Sec. 3.5, Table S2 |
|  | Hardware & software environment | PyTorch, Scikit-learn, XGBoost, Life-lines; reproducible execution logs in metrics.json | Sec. 3.5, Sec. S2 |
| **4. Model Evaluation** | Data partitioning & leak-free design | 5-fold cross-validation; pretraining en-coders strictly isolated from downstream test splits | Sec. 3.5, Sec. S2 |
|  | Baseline models & fairness | Identical splits and preprocessing for Raw features, PCA, scVI, FlowSOM, and Ran-dom Forest | Table 3, Table 5, Table 6 |
|  | Discrimination & calibration metrics | Macro/Weighted F1, ROC AUC, Accuracy, Silhouette width, Effect size (*σ*), ARI, NMI | Tables 3 to 6, Figs. 2 to 5 |
|  | Survival analysis metrics | Harrell's *C*-index and log-rank test *p*-values on high- vs. low-risk Kaplan-Meier stratifi-cation | Table 6, Fig. 6 |
|  | Variance & confidence intervals | 5-fold cross-validation standard deviations reported for all test splits | Tables 3 to 6, Table S3 |
| **5. Explainability & Interpretability** | Manifold geometry & biological fidelity | 2D UMAP projections, per-class silhouette analysis, Hungarian-matched clustering | Fig. 3, Table S8 |
|  | Clinical gating tree concordance | Stage-specific bi-axial gating back-projection (88 to 98.5% concordance in SupCon) | Fig. 4, Fig. S3 |
|  | Error analysis & limitations | Inductive bias divergence (decision trees vs. manifolds); sample size limitations in sur-vival | Sec. 5.2.1, Sec. 5.2 |
|  | Code & data availability | Public dataset accessions cited; proprietary source code shared with editors and review-ers for peer review | Backmatter, Sec. S2 |

**Table S2.** VICReg self-supervised encoder hyperparameters and loss coefficients by benchmark cohort.^a^.

| Cohort | Hidden Dim | Masking Rate *p* | Noise σ | (λ, μ, ν) |
|---|---|---|---|---|
| Levine-13 | 256 | 0.00 | 0.03 | (25, 25, 5) |
| FlowCAP-II | 128 | 0.15 | 0.30 | (25, 25, 1) |
| FlowCAP-IV | 128 | 0.08 | 0.05 | (25, 25, 1) |
| OMIP-030 (Tier 2) | 128 | 0.15 | 0.15 | (25, 25, 1) |

<sup>a</sup>These are the self-supervised VICReg encoder’s own hyperparameters (embedding dim 16, projection dim 32, 3 residual blocks throughout, AdamW). The separate directly-supervised SupCon classifier used for OMIP-030’s classification ceiling (Section 4.2.4) has its own, independently-tuned configuration: hidden dim 256, mask\_prob 0.05, noise σ 0.02, temperature 0.1, 40 epochs, AdamW (lr 1 × 10<sup>−3</sup>), CosineAnnealingLR; it is not a VICReg model and is not covered by this table.

**Table S3.** Levine-13, identical 5-fold XGBoost-probe CV for every representation row. The supervised rows represent a ceiling reference, not a competing self-supervised method.

| Method | Accuracy | Macro F1 |
|---|---|---|
| *Supervised upper bound (trained directly on labels, raw features)* |  |  |
| Random Forest | 0.998 ± 0.000 | **0.945** ± **0.004** |
| XGBoost | 0.996 ± 0.001 | 0.929 ± 0.006 |
| *Self-supervised / unsupervised representations (probe-evaluated)* |  |  |
| PCA | 0.977 ± 0.001 | 0.847 ± 0.013 |
| scVI | 0.966 ± 0.001 | 0.811 ± 0.010 |
| VICReg (encoder only) | 0.965 ± 0.001 | 0.812 ± 0.011 |
| VICReg (PCA-anchored) | 0.976 ± 0.001 | 0.843 ± 0.014 |
| FlowSOM (24 clusters) | 0.632 ± 0.117 | 0.306 ± 0.117 |

**Table S4.** Levine-13 credit decomposition: what does the self-supervised training regime itself contribute, isolated from the frozen linear component?

| Component | Macro F1 |
|---|---|
| PCA baseline (independently fit, standard protocol) | 0.847 ± 0.013 |
| Frozen PCA skip alone (sanity check, same pretraining pool) | 0.840 ± 0.010 |
| + trainable VICReg residual (self-supervised contribution) | 0.843 ± 0.014 |
| *Isolated marginal contribution of VICReg training* | +0.003 |

**Table S5.**
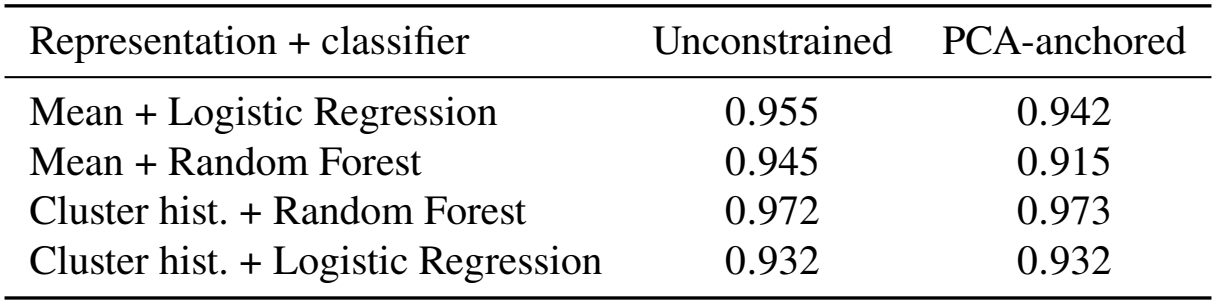
FlowCAP-II with vs. without PCA-anchoring. Unlike Levine-13, PCA is a weaker baseline than unconstrained VICReg on this cohort (0.926, Table 5), and anchoring does not help.

| Representation + classifier | Unconstrained | PCA-anchored |
|---|---|---|
| Mean + Logistic Regression | 0.955 | 0.942 |
| Mean + Random Forest | 0.945 | 0.915 |
| Cluster hist. + Random Forest | 0.972 | 0.973 |
| Cluster hist. + Logistic Regression | 0.932 | 0.932 |

a. **Leukemic Myeloblasts (Cluster 3)**: Contains 592,433 cells, of which 70.18% are derived from AML patients (a 5.82× fold enrichment above the cohort baseline of 12.05%). Morphologically and phenotypically, Cluster 3 exhibits the hallmark myeloblast profile: intermediate forward scatter (FSC), low side scatter (SSC), CD45^dim^ expression, and high co-expression of immature stem and myeloid progenitor markers (CD34^+^, CD117^+^, HLA-DR^+^, CD38^+^).
b. **Monocytic / Precursor Lineages (Clusters 12 and 14)**: Cluster 14 (25.82% AML cells) captures an activated HLA-DR^+^CD38^+^ monocytic precursor population, while Cluster 12 isolates large activated monocyte-like cells (FSC^high^CD45^high^).
c. **Non-Malignant Mature Subsets (Clusters 2, 7, 9)**: These clusters exhibit high CD45 expression, low side scatter, and negligible immature markers (CD34^−^CD117^−^), corresponding to mature lymphocytes (T, B, and NK cells) with low AML cell fractions (12.8–14.7%, tracking normal background).
d. **Mature Granulocytes (Clusters 4, 6, 8, 10)**: Characterized by prominent high side scatter (SSC^high^) and intermediate CD45, these clusters represent mature neutrophils and granulocytes, exhibiting low AML cell fractions (1.4–4.4%).

This demonstrates that Elflow’s patient-level classification is directly rooted in genuine biological discrimination: Random Forest classifiers heavily weight the frequency of Cluster 3 to distinguish AML patients from healthy donors without requiring manual gating boundaries.

**Figure S1.**
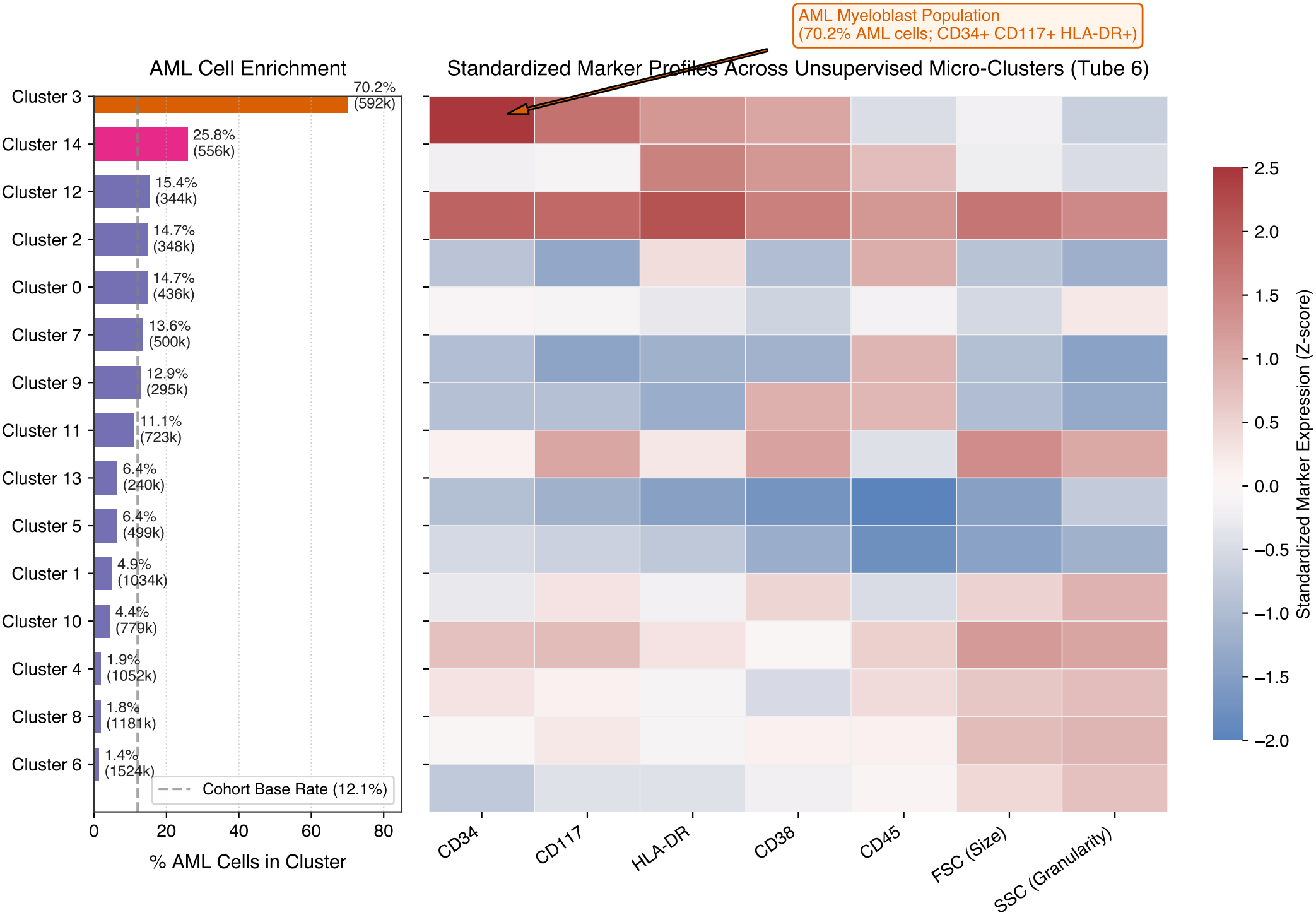
FlowCAP-II unsupervised micro-cluster profiling and phenotypic interpretation (Tube 6). Left: Percentage of AML patient cells landing in each of the 15 unsupervised micro-clusters (ordered by AML enrichment; total event count in parentheses). The vertical dashed line indicates the overall cohort baseline AML prevalence (12.05%). Right: Standardized marker expression heatmap (*Z*-scores) across the 7 parameters in Tube 6 (CD34, CD117, HLA-DR, CD38, CD45, FSC, SSC). Cluster 3 (orange highlight) isolates the primary leukemic myeloblast population (70.18% AML cells, *>* 5.8× cohort enrichment), characterized by classic CD34^+^CD117^+^HLA-DR^+^CD38^+^CD45^dim^SSC^low^ co-expression.

**Table S6.**
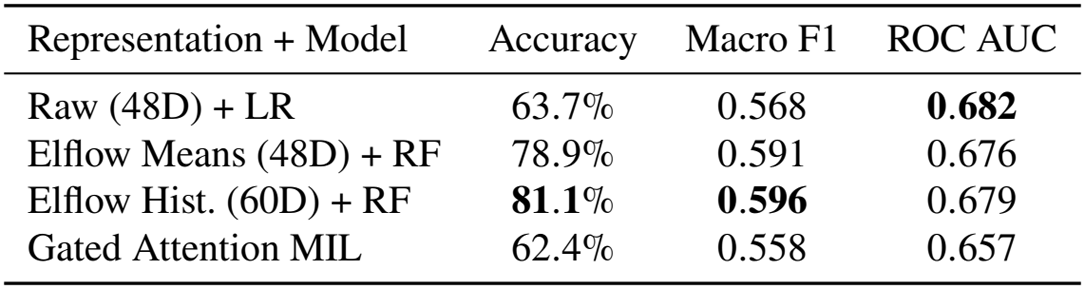
FlowCAP-IV binary progression classification across patient-level representations (5-fold CV).

| Representation + Model | Accuracy | Macro F1 | ROC AUC |
|---|---|---|---|
| Raw (48D) + LR | 63.7% | 0.568 | **0.682** |
| Elflow Means (48D) + RF | 78.9% | 0.591 | 0.676 |
| Elflow Hist. (60D) + RF | **81.1**% | **0.596** | 0.679 |
| Gated Attention MIL | 62.4% | 0.558 | 0.657 |

### S6. FlowCAP-IV: Full Classification Results

Figure S2 shows the complete evaluation grid: 9 patient representations (raw, PCA, and Elflow, each in concatenated and differential forms, plus the Elflow histogram and hybrid representations) evaluated with Logistic Regression, Random Forest, and XGBoost under identical 5-fold cross-validation. In this benchmark, each subject is profiled across three experimental tubes (antigen-stimulated Env, Gag, and unstimulated control) measuring 16 markers each; concatenated raw and mean-pooled representations therefore yield 3 × 16 = 48D feature vectors, concatenated micro-cluster histograms yield 3 × 20 = 60D vectors, and the hybrid representation combines both into 48D + 60D = 108D.

### S7. OMIP-030: Quality Control and Fine Phenotypic Resolution Details

#### S7.1. Quality Control Ground-Truth Benchmark and Automated Segmentation

On the 22-channel OMIP-030 dataset (1,664,738 raw events), ground-truth reference targets for quality control benchmarking were established using the published OMIP-030 manual gating protocol based on morphological scatter and viability criteria:

**Figure S2.**
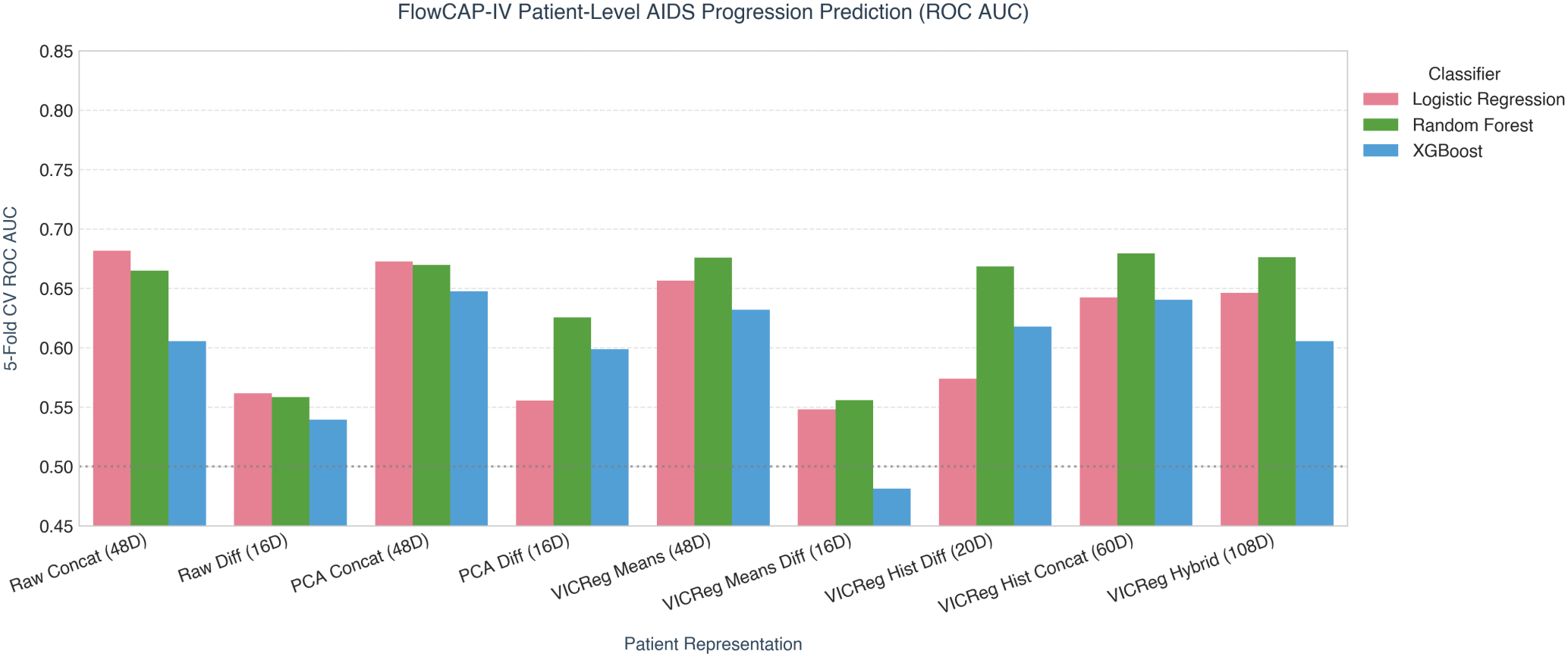
FlowCAP-IV progression-classification ROC AUC, all 9 representations × 3 classifiers, 5-fold CV.

a. **Debris**: FSC-A *<* 35,000 or SSC-A *<* 5,000 (122,948 events, 7.39%).
b. **Doublets / Aggregates**: Non-linear pulse width events satisfying FSC-W ∉ [62k, 85k] or SSC-W ∉ [58k, 82k] (208,465 events, 12.52%).
c. **Non-Viable Cells**: Live/Dead Blue^+^ events (*>* 4,000 counts) within the singlet gate (174,383 events, 10.47%).
d. **Clean Live Singlets**: 1,158,942 events (69.62% yield).

To eliminate manual gating during pipeline execution, an automated 4-class gradient-boosted decision tree (XGBoost) within Elflow was trained on morphological scatter (FSC, SSC) and viability channels to autonomously segment raw events into clean live singlets, doublets, dead cells, and debris. Evaluated on a held-out test split (20% split, *n* = 332,948 events), the automated model achieves Macro F1 0.9942 and overall accuracy 99.69% (Table S7), confirming that cytometric noise can be autonomously filtered at inference time without human intervention prior to downstream representation learning.

**Table S7.** OMIP-030, automated 4-class quality control segmentation performance on held-out test split (20% split, *n* = 332,948 events).

| Class | Precision | Recall | F1-Score | Event Count |
|---|---|---|---|---|
| Clean Live Singlets | 0.998 | 0.997 | 0.998 | 231,788 |
| Doublets | 0.992 | 0.994 | 0.993 | 41,693 |
| Dead Cells | 0.995 | 0.996 | 0.996 | 34,877 |
| Debris | 0.990 | 0.989 | 0.989 | 24,590 |
| **Macro Average** | **0.994** | **0.994** | **0.994** | **Accuracy: 99.69%** |

**Table S8.** OMIP-030, per-class cluster cohesion (*s*_*i*_) and neighborhood purity (*k* = 15) for raw markers vs. Cascaded Elflow embeddings, evaluated on the full *N* = 9,949 expert-gated CD4^+^ cohort (FR-FCM-ZZWU).

| Biological Subset | Events | Raw *k*NN Purity | Elflow *k*NN Purity | Silhouette (*s<sub>i</sub>*), Raw → Elflow |
|---|---|---|---|---|
| Naive | 1,630 | 52.8% | 54.5% | +0.050 → +0.007 |
| Central Memory | 369 | 18.0% | 17.0% | −0.006 → −0.059 |
| Effector Memory | 1,871 | 43.6% | 40.8% | −0.100 → −0.155 |
| TEMRA | 3,476 | 64.0% | 61.7% | −0.045 → −0.074 |
| Tfh | 651 | 27.1% | 15.6% | −0.017 → −0.090 |
| Th1 | 1,141 | 39.0% | 34.4% | −0.041 → −0.027 |
| Th2 | 181 | 7.7% | 7.3% | −0.081 → −0.093 |
| Th17 | 162 | 21.0% | 14.4% | **+0.158 → +0.225** |
| T<sub>regs</sub> | 468 | 27.0% | 28.6% | **−0.040 → +0.048** |
| **Overall** | **9,949** | **47.9%** | **45.5%** | −0.033 → −0.061 |

Under the alternative, naive population-std normalization (rather than the pooled within-class-std effect size reported in Table 3), mean pairwise centroid distance across all 36 subset pairs tells a similar story: Raw 1.881, PCA 1.778 (−5.5% vs. Raw), scVI 2.239 (+19.0%), Flat VICReg 2.748 (+46.1%), Cascaded Elflow 2.687 (+42.8%); Flat again expands separation furthest, with Cascaded close behind, under either normalization.

As shown in Table S8, the global silhouette score is negative for both raw markers (−0.033) and Cascaded Elflow (−0.061): neither representation produces well-separated clusters by this metric on the full 9-class population, and Cascaded Elflow is directionally worse, not better, in aggregate. Local neighborhood fidelity (*k*NN purity, *k* = 15) is similarly slightly lower under Cascaded Elflow overall (45.5% vs. 47.9% raw). Only 2 of the 9 subsets show a genuine silhouette gain under Cascaded Elflow: Th17 (+0.158 → +0.225) and T_regs_ (−0.040 → +0.048, moving from slightly negative to clearly positive), a narrow silhouette-only improvement, not a broader clustering-quality win (Table 3). The remaining 7 subsets are less cohesive under Cascaded Elflow than under raw markers on both metrics.

#### S7.2. Lineage-Dominance Ablation: A Methodologically Cleaner Test of the Cascade Mechanism

The main-text benchmark (Table 3) compares Flat and Cascaded VICReg on the same CD4^+^-restricted pretraining pool (23,115 cells), which has a real limitation: a lineage-homogeneous pool contains no genuine multi-lineage variance for a flat encoder to be dominated by, so this comparison may understate how severely a flat encoder collapses under real lineage dominance. We therefore separately trained Flat VICReg on the genuinely multi-lineage Live population (61,229 cells spanning CD4^+^/CD8^+^ T cells, B cells, and other major lineages, gated directly from the real FlowJo workspace, with test-labeled cells excluded from pretraining exactly as in the main benchmark) and compared it to Cascaded VICReg trained on the standard CD4^+^-restricted pool, using the identical real ContrastiveEncoder/VICRegTrainer architecture and hyperparameters (Supplementary Table S2) as the main benchmark.

**Table S9.** Lineage-dominance ablation: Flat VICReg pretrained on the genuinely multi-lineage Live population (61,229 cells) vs. Cascaded VICReg on the standard CD4^+^-restricted pool (23,115 cells).^a^.

| Metric | Flat VICReg (multi-lineage) | Cascaded VICReg |
|---|---|---|
| Effective rank | 5.65/16 | **7.53/16** (+33.4%) |
| Tree Probe Macro F1 | 0.346 | **0.444** |

<sup>a</sup>Unlike the main-text comparison (Table 3), where both Flat and Cascaded train on the same CD4<sup>+</sup>-restricted pool, here Flat is deliberately pretrained on a pool with genuine lineage variance to dominate, isolating the cascade mechanism more cleanly. These two numbers were captured from this run's console log rather than a saved metrics.json artifact, unlike every other number in this paper; we report them for completeness but flag the difference in provenance rigor explicitly.

The relative effective-rank gain is close to the main-text, matched-data comparison (+33.4% here vs. +16.6% there), and Tree Probe F1 also improves under cascading in both constructions, indicating the cascade mechanism’s benefit is not an artifact of either the training code path or the specific pretraining pool used.

#### S7.3. Label-Free Anchor-Guided Cluster Profiling: The Deployment-Realistic Unsupervised Evaluation

The clustering-quality metrics above (Table 3) use a generic *K*=9 Hungarian-matched assignment, which is standard practice for comparing representation quality but does not reflect how an unsupervised pipeline would actually be deployed (an arbitrary cluster count, no confidence filtering). To evaluate each representation under a more realistic unsupervised deployment scenario, we separately cluster each representation directly (MiniBatchKMeans, *K* ∈ {9, 15, 20, 30, 50}) and label each cluster via the anchor-guided profiling procedure of Section 3.4.1, with anchors defined in the raw 10D subset-marker space (rather than the embedding space itself) specifically for this sweep, so that the effect of cluster count *K* can be isolated from the effect of anchor definition.

**Table S10.** Label-free anchor-guided cluster profiling: full-population F1 (unlabeled clusters scored as wrong) by cluster count *K*.

| Representation | *K*=9 | *K*=15 | *K*=20 | *K*=30 | *K*=50 |
|---|---|---|---|---|---|
| Raw | 0.113 | 0.158 | 0.159 | 0.186 | 0.205 |
| PCA | 0.100 | 0.125 | 0.186 | 0.187 | 0.197 |
| scVI | 0.130 | 0.151 | 0.144 | 0.177 | 0.177 |
| Flat VICReg | **0.174** | **0.195** | **0.241** | **0.241** | **0.269** |
| Cascaded Elflow | 0.139 | 0.158 | 0.193 | 0.199 | 0.217 |

Flat VICReg wins at every *K* tested (+31% relative to Raw at *K*=50); Cascaded Elflow also beats Raw consistently, by a smaller margin (+6% at *K* = 50). Coverage (fraction of cells landing in a cluster with at least one anchor) tells a complementary story: both VICReg variants hold complete, 100% coverage at every *K* from 9 to 50, while Raw drops to 96.3%, PCA to 97.9%, and scVI to 94.4% coverage at *K*=50: self-supervised representations are simultaneously more accurate *and* more robust to leaving cells unclustered in this deployment scenario, even though the flat (not cascaded) variant is the stronger of the two here. This raw-space-anchor sweep is what motivated using a much larger cluster count for the main-text result: a parallel sweep with anchors defined in the embedding space itself (Section 3.4.1) showed the same resolution problem on the Cascaded Elflow embedding specifically. *K*=30 (the resolution used in an earlier pass of this analysis) misses Th17, Th2, and Tfh entirely (zero predicted cells, despite Th17 having the best per-class silhouette of all 9 classes in this embedding, a pure clustering-resolution problem, not a labeling failure); *K*=50 recovers Th17, *K*=75 recovers Tfh, and *K*=150 is near-optimal; Th2 remains unrecovered even at *K*=200, consistent with its negative per-class silhouette (−0.093, Table S8): no clustering resolution fixes a class the representation itself does not distinguish. Embedding-space anchors at *K*=150 is therefore the configuration used to produce the Cascaded Elflow column of Figure 4 (Section 3.4.1), not the raw-space-anchor configuration tabulated above, which was retained at its original, lower *K* range specifically to isolate the anchor-definition question from the resolution question.

#### S7.4. Canonical 4-Stage Hierarchical Gating Tree

To resolve all 9 canonical CD4^+^ T-cell subpopulations without loss of high-dimensional context, manual flow cytometric analysis in OMIP-030 employs a sequential 4-stage bi-axial gating hierarchy (Figure S3). Starting from clean live CD4^+^ singlets (*N* = 9,949 reference events across all 9 canonical subsets, used throughout Table 3 and Table S8):

a. **Stage 1 (Memory Differentiation, Row 1)**: Resolves four discrete maturation quadrants on the CCR7 × CD38 plane: Naive (CCR7^+^CD38^+^), T_cm_ (CCR7^−^CD38^+^), T_em_ (CCR7^−^CD38^−^), and TEMRA (CCR7^+^CD38^−^). This is corrected from the textbook CD45RA × CCR7 pairing: tracing the exact RectangleGate boundaries in this sample’s own FlowJo workspace showed CD45RA is never the restrictive dimension anywhere in this stage’s hierarchy (both parent branches span its full acquisition range), while CCR7 (parent gate) then CD38 (leaf gate) form a clean, shared-boundary split; confirmed empirically on real labeled data (per-class arcsinh medians cleanly separate on both thresholds).
b. **Stage 2 (Regulatory T-Cells, Row 2)**: Directly isolates the CD25^high^CD127^low^ T_reg_ lineage from the parent CD4^+^ compartment; this plane matches the reference leaf gate’s own name exactly and was not revised.
c. **Stage 3 (Type 1 vs. Type 17 Helper Polarization, Row 3)**: Restricts to the CD45RA^−^ antigen-experienced memory pool and segregates Th1 from Th17 on CCR4 × CCR10, corrected from the textbook CCR6 × CXCR3 pairing after tracing the real gate hierarchy (Th1 and Th17 share a CCR6^−^ grandparent gate, then diverge on CCR4 and CCR10, not CCR6/CXCR3); confirmed empirically (Cohen’s *d*: CCR4 2.08, CCR10 1.77, vs. CCR6 1.22, CXCR3 0.44 for the original pair).
d. **Stage 4 (Type 2 vs. Follicular Helper Polarization, Row 4)**: Restricts to the CD45RA^−^ memory pool and segregates Th2 (CCR4^+^CXCR5^−^) from Tfh (CXCR5^+^CCR4^−^) on CCR4 × CXCR5; independently confirmed as the empirically best-separating pair for this split (Cohen’s *d*: CCR4 1.49, CXCR5 1.37, both ahead of alternatives) and not revised.

This hierarchical structure establishes the ground-truth benchmark against which the 2D bi-axial gating concordance of self-supervised representations is evaluated in Main Text Figure 4.

**Figure S3.**
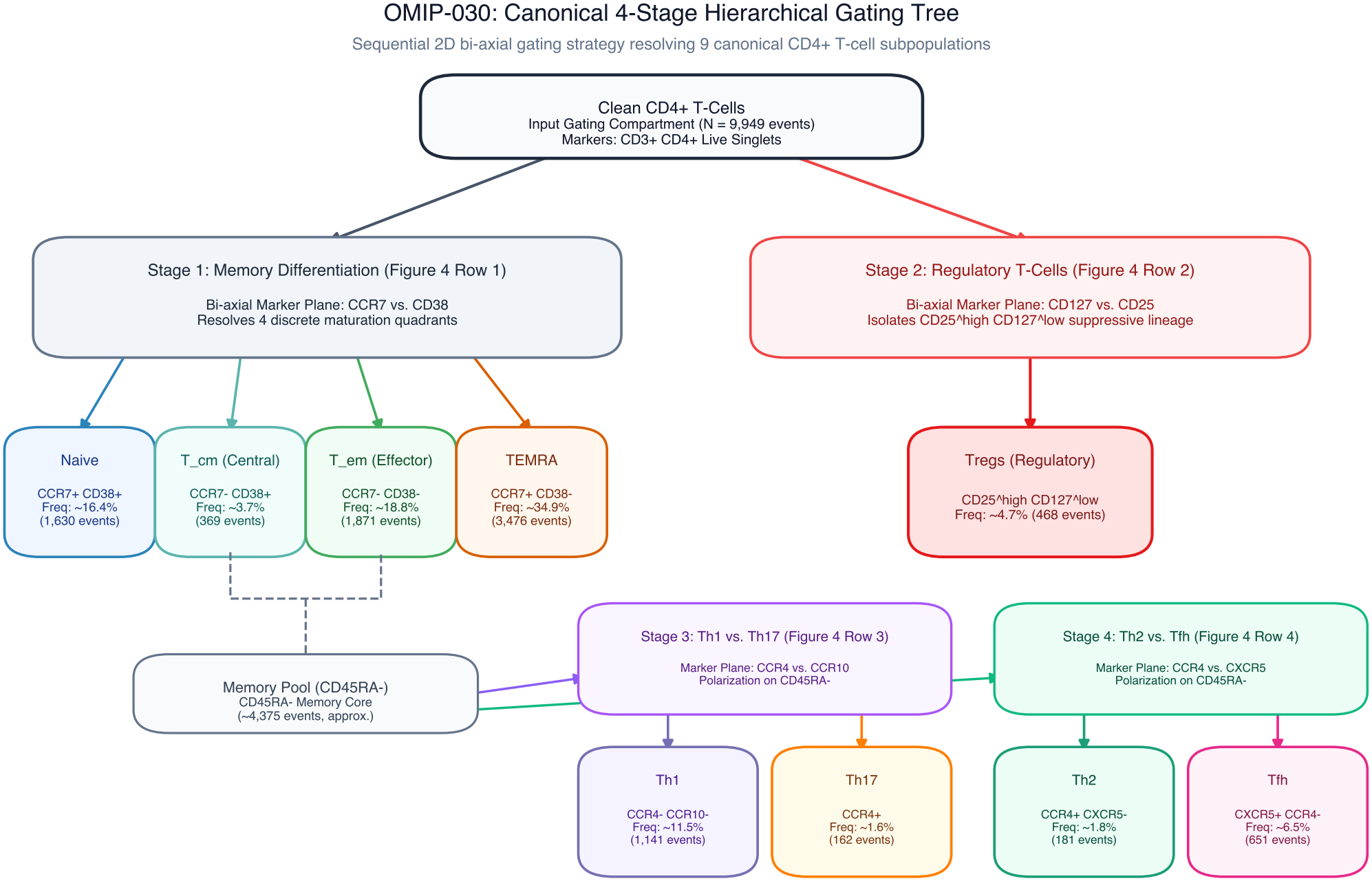
OMIP-030 canonical 4-stage hierarchical gating tree. Flowchart showing the sequential 2D bi-axial gating strategy used to isolate all 9 reference CD4^+^ T-cell subpopulations from clean live singlets (*N* = 9,949 events). Each branching stage directly corresponds to the respective row (Stages 1 to 4) evaluated for gating concordance in Main Text Figure 4.

## Notes

### Competing Interest Statement

The authors have declared no competing interest.

